# Conservation of the hydrogen-bond network in bacterial response regulators

**DOI:** 10.1101/2025.09.13.674700

**Authors:** Maham Hamid, Ishrat Jabeen, Safee Ullah Chaudhary, Shahid Khan

## Abstract

The bacterial response regulator (RR) superfamily is activated by single aspartyl phosphorylation to modulate a distant target binding surface for diverse functions. The enteric CheY RRs, which represent the chemotaxis subfamily, have been extensively characterized. Their native, chemical or genetically-altered crystal structures have revealed an essential role for water-mediated hydrogen bonds (H-bonds) in activation. Here, we use molecular dynamics (MD) to compare the protein-water H-bond network in basal and *in-silico* phosphorylated conformations. We supplement the MD with energy frustration profiles for atomic structures and models from selected RR superfamily representatives. The energetically frustrated phosphorylation pocket consists of the conserved aspartate triad for phosphorylation, plus associated structural waters and residues for Mg^2+^ ion coordination. It orchestrates the H-bond network characterized here in atomic detail. The network has an energetically stable core. It’s plastic nodes switch bonding states coupled to loop flexibility and sidechain rotations. Mutual information reveals that the long-range, dynamic networks respond to single H-bond transitions. The network centrality of the phosphorylation pocket, connected to the target binding surface by water-mediated channels via the conserved switch residues (T87, K109), increases upon phosphorylation. Analysis of other RR representatives suggests this design is a generic feature of RR allostery with subtle, function-dependent differences. The water contribution may prove critical for the design of specific RR sub-family specific, allosteric inhibitors.

## Introduction

The bacterial response regulator (RR) superfamily of sensory proteins mediates diverse responses that dictate how bacteria interact with their environment (Gao et al., 2019). Most members bind DNA and, to a lesser extent, protein and RNA for functions that include gene regulation, metabolism, phosphate regulation, osmosensing and chemotaxis (Gao et al., 2007). In the years since the first structures of the canonical enteric *E. coli* and *S. typhimurium* CheY chemotaxis proteins were published (Stock et al., 1989; Volz and Matsumura, 1991), over 2 million sequences and 100 near-atomic crystal structures of superfamily members have been deposited in protein databases, making the superfamily the ninth most abundant in nature (Kennedy et al., 2022).

Signal transduction in this family involves phosphorylation of a conserved aspartyl residue, as first demonstrated for the enteric CheY chemotaxis proteins (Bourret et al., 1990; Lukat et al., 1991). Henceforth, “CheY_es_*“* will refer jointly to the *E. coli / S. typhimurium* CheY chemotaxis proteins (>98% sequence identity), with CheY_es_ numbering applied to residue positions, unless otherwise specified. The transition from the basal (inactive) to phosphorylated (active) conformation is best understood for CheY_es_. Crystal structures have revealed the essential role of magnesium (Mg^2+^) as a cofactor (Stock et al., 1993). The conformational changes triggered upon activation (Cho et al., 2000; Lee et al., 2001) by phosphorylation of a single aspartate (D57) by a cognate kinase elicit changes at a common target binding surface (Zhu et al., 1997b). The D57 phosphoaspartyl linkage is labile, so beryllium fluoride (BeF_3_), a phosphoaspartyl analog, is commonly used to mimic the activated state (Petsko, 2000). A library of CheY crystal structures with residue substitutions has linked conformational state to chemotactic function (Table S2 (Wheatley et al., 2020)) mediated by two residues (T87, Y106). Residue substitutions in T87 uncouple phosphorylation from chemotaxis (Ganguli et al., 1995; Jiang et al., 1997). Y106 sidechain rotation correlates with chemotactic response (Zhu et al., 1997a). Acetylation of the conserved K109, and K91, enhances motor CW rotation analogous to phosphorylation (Fraiberg et al., 2015). Database-guided mutagenesis (Kennedy et al., 2022) has revealed that residue determinants of RR phosphorylation/dephosphorylation kinetics vary by 5 or more orders of magnitude between family members (Immormino et al., 2016; Page et al., 2016). These kinetics are rapid for CheY_es_ (Foster et al., 2021; Straughn et al., 2020), consistent with the sub-second chemotactic excitation and adaptation kinetics (Khan et al., 1995). However, other RRs have important differences in the downstream reactions mediated by aspartyl phosphorylation. For example, phosphorylation facilitates homodimerization in the transcriptional regulator OmpR/PhoB family to increase DNA affinity (Toro-Roman et al., 2005).

Water is an integral component of protein H-bonded networks. The role of bound waters, long-recognized as important for protein folding (Papoian et al., 2004), is increasingly appreciated as critical for allosteric communication (Bondar, 2022). For CheY_es_, dynamic H-bonds are central to allosteric coupling (Dyer and Dahlquist, 2006) and references therein, as for many protein families (Fersht et al., 1985). Protein-water H-bond dynamics have not been considered in molecular dynamics (MD) studies on CheY ((Foster et al., 2021; Foster and West, 2017; Wheatley et al., 2020) and references therein) or other RRs that have analyzed C^α^ residue-residue couplings.

The X-ray footprinting data on sidechain solvent accessibility and the C^a^-based community networks introduced in our work (Wheatley et al., 2020) reported that subtle sidechain rotamer selection and loop immobilization drove CheY activation. These changes altered the solvation of the solvation site (Hamid et al., 2022), consistent with the analysis of the static crystal structures by the Bridge2 (Siemers and Bondar, 2021) and Frustratometer2 (Parra et al., 2016) algorithms. However, a D13KY106W double-mutant to mimic activation was used in these studies, preventing a mechanistic analysis of the phosphorylation pocket. We have now developed methodology to characterize the basal (Mg^2+^) and phosphorylated (Mg^2+^/D57_PO4_) CheY conformational landscapes *in silico* (Hamid et al., 2025). Here we exploit this methodology for simulation of CheY_es_ basal (2CHE.pdb (Stock et al., 1993)) and phosphorylated conformations (1FQW_PO4_), the latter generated from BeF_3-_activated 1FQW.pdb (Lee et al., 2001). Protein-water H-bond networks were then constructed for selected representatives of the RR superfamily and compared to these template networks. Their subsequent analysis revealed the central role of water in the architectural dynamics of the phosphorylation pocket and the allosteric relay linking it to the target binding surface.

## Results

### 1. Validation and dynamics of *in-silico* phosphorylated CheY

We first confirmed that our dianionic phosphate modified structure (1FQW_PO4_) reported the activated-mediated changes of the angles and bond distances visualized in BeF_3_-modified CheY_es_ (1FQW), a mimic of the phosphorylated active state (Lee et al., 2001) (**Fig. 1A**). See also (Foster and West, 2017). There are five conserved residues (D12, D13, D57, K109, T87) in the RR superfamily (Gao et al., 2019). D57 phosphorylation requires Mg^2+^ coordination with D12, D13 (Stock et al., 1993), forming the “phosphorylation pocket”. K109 forms an activation-dependent salt-bridge with D57, while T87 mediates conformational coupling between the D57 phosphorylation site and Y106 at the target binding surface(Dyer et al., 2004). The Y106 is substituted, predominantly by other aromatic residues, in other RR superfamily members, but its internal rotation, seenin BeF_3_-modified CheY_es_ has been a diagnostic of CheY_es_ activation. 1FQW_PO4_ Y106 and T87 had an inward orientation relative to the basal form (2CHE). In addition, D57 bonded with K109, T87 and N59, while Y106 bonded with E89, all as observed in 1FQW, We carried out simulations to record the variations in these parameters (**Fig. 1B**) and the couplings between them as reported by the normalized mutual information (nMI) (**Fig. 1C**). The differences between the basal and phosphorylated states were maintained. The couplings were not absolute, but significant. The largest difference between the states was in the T87-D57 distance due to bond formation upon phosphorylation. T87-D57 bond formation resulted in the most pronounced increase in the dynamic couplings mediated by phosphorylation.

**Fig. 1.**
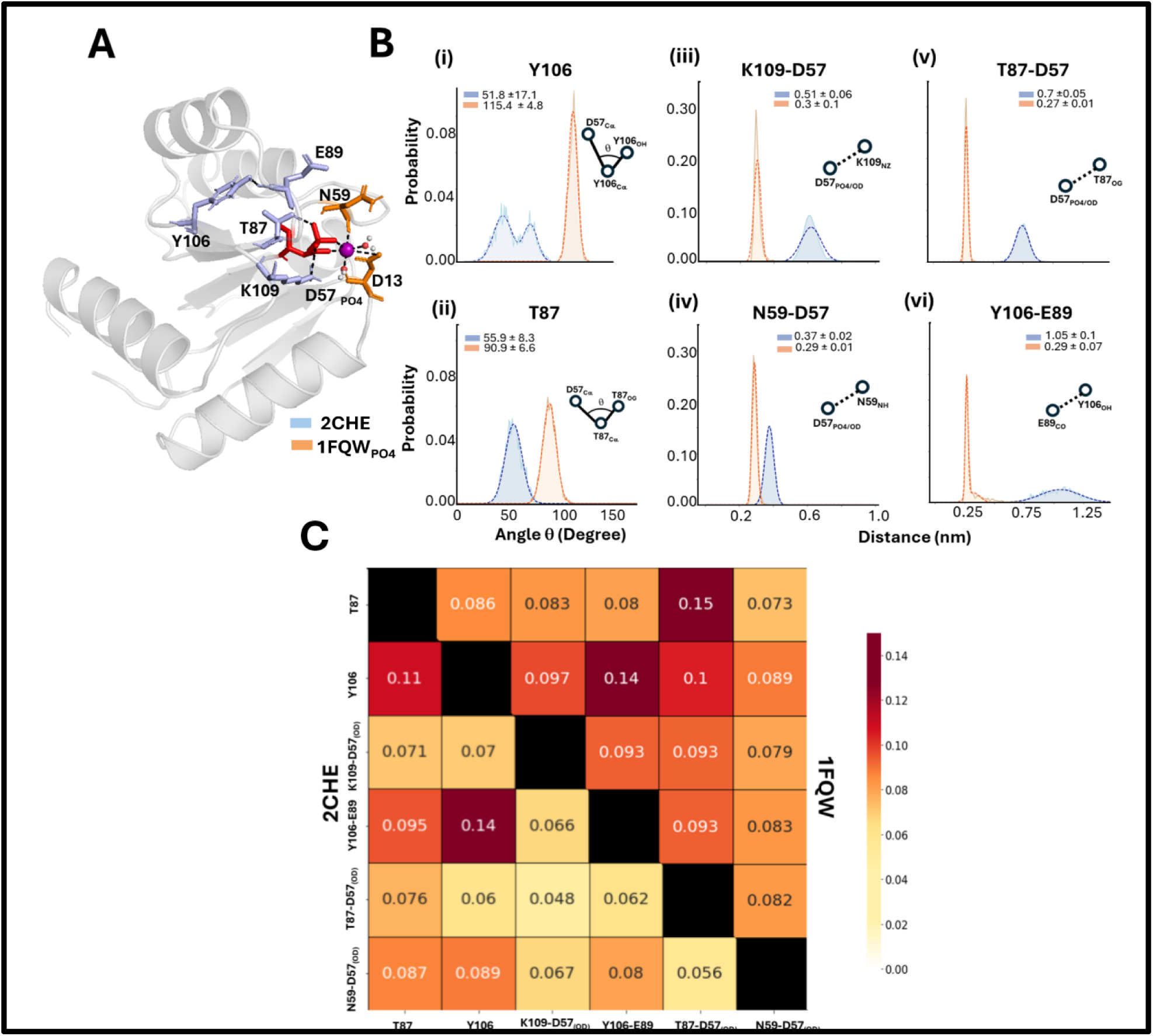
D57 phosphoaspartyl activation tracked by indicators identified from the BeF_3_-activated CheY structure. **A.** Cartoon representation of the crystal structure (1FQW.pdb), with the D57 phosphoaspartyl modification (red). D57_PO4_, like BeF_3,_ contributes to the Mg^2+^ ion (magenta) coordination to replace a bound water molecule. **B.** The key changes triggered by activation are internal rotations of the **(i)** Y106 and **(ii)** T87 sidechains,andbondformationbetween **(iii)** K109-D57, **(iv)** N59-D57. **(iv)** T87-D57 and **(vi)** Y106-E89**. C.** The dynamic couplings between these indicators are measured with the nMI heatmap.

### 2. Fold variation in the CheY superfamily

The crystal structures of the chemotactic mutants have not shown a clear correlation between behavioral phenotype and the CheY_es_ fold. For example, the constitutively active D13KY106W double mutant is more similar to basal CheY_es_ than BeF_3_-modified CheY (Dyer et al., 2004). Therefore, we explored more sensitive measures to relate fold to function – by simulation as well as multi-step clustering of the available databases.

First, we ran long (1μs) simulations of the 2CHE and 1FQW_PO4_ structures as well as the structure of the Mg^2+^-minus apo-CheY(3CHY)(Volz and Matsumura, 1991). The variations between replicates were assessed with PCI-PC2 plots (**Fig. S1**). The relatively isotropic 3CHY PC1-PC2 plot became more anisotropic in 2CHE and 1FQW_PO4_ as Mg^2+^ chelation allowed CheY to access more diverse conformational states. Next, we concatenated the 2CHE, 3CHY, and 1FQW_PO4_ replicate trajectories. The PC1-PC2 plot of the concatenated 2CHE and 1FQW_PO4_ trajectories revealed that overlap was restricted to a distinct C^α^ fold subspace (**Fig. 2A**). Cluster analysis of the concatenated trajectory identified 7 distinct clusters comprising 2CHE alone (4), 1FQW_PO4_ alone (3), and mixed 2CHE-1FQW_PO4_ (1). Chelation affects the CheY fold since apo-CheY (3CHY) formed a distinct cluster (**Fig. 2B**). The analysis reinforced previous conclusions that CheY undergoes conformational selection upon phosphorylation.

**Fig. 2.**
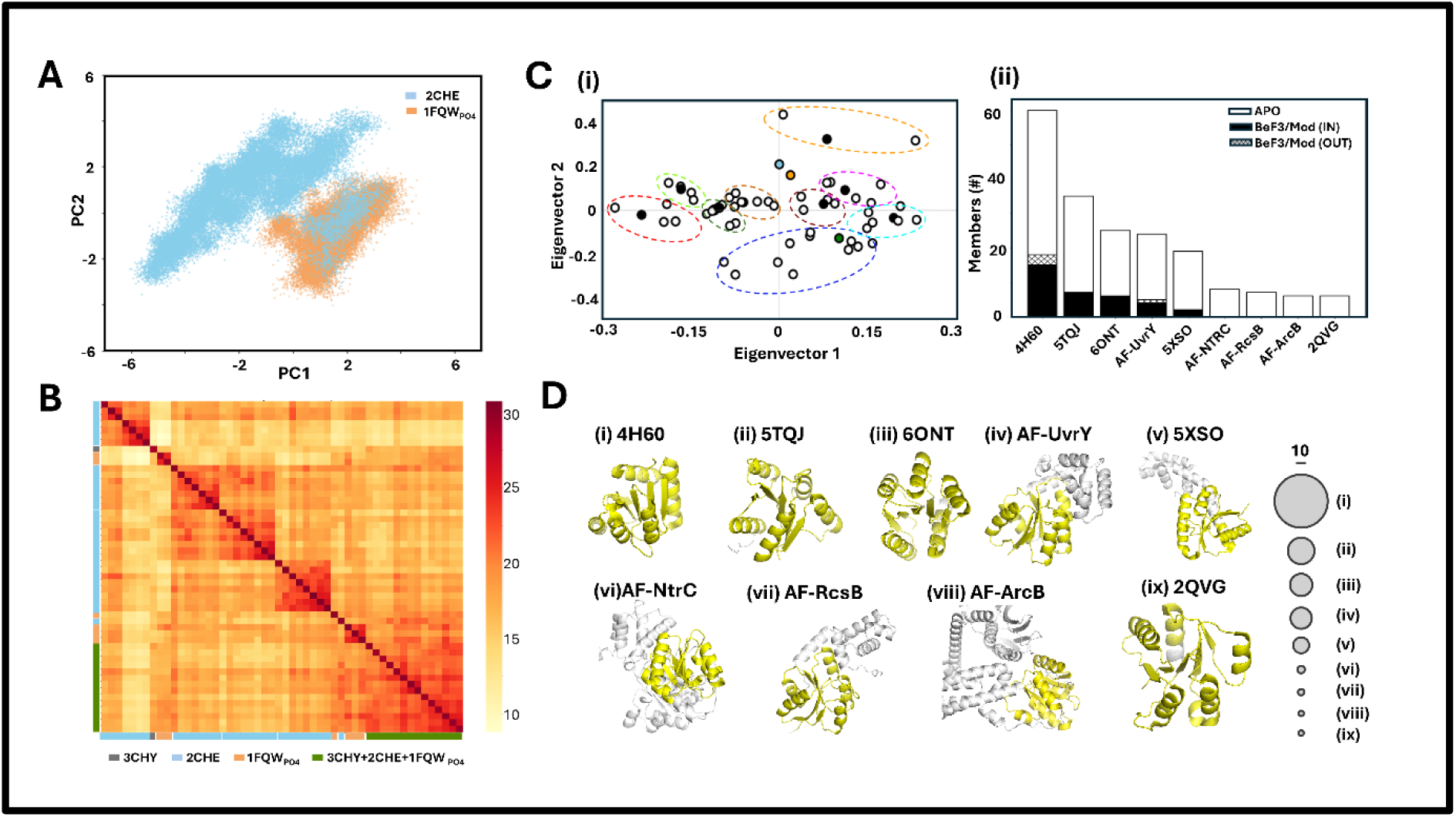
A, B Fold variations between basal (2CHE) and active (1FQWPO4) states. 1μs simulations (3 replicates)**. A.** PCI-PC2 plot comparison of the basal and activated CheY backbone (C^a^) conformational space. **B.** DALI heatmap reports homogeneous 2CHE (cyan). 1FQW_PO4_ (orange) and mixed (2CHE + 1FQW_PO4_) group conformations. **C. (i)** The DALI correspondence plot of the 59 representative structures obtained from CD-Hit. **(ii).** Distribution of the total (196) RR structures partitioned among the final 9 representatives after reduction of the post CD-Hit structures by DALI fold correspondence and CATH-based functional overlap. The size of the IN (black) and OUT (bar pattern) Y106 (or equivalent) rotamer position fractions in structures activated by BeF_3_ /other (black) chemical modification or mutation. **D.** The 3D structures of the final cluster representatives. The RR domains are colored yellow.

Next, we examined RR superfamily sequence and structure databases. AlphaFold (AF) models of entries in the seed PF00072 multiple sequence alignment (MSA), where available, were downloaded from InterPro or constructed *de novo*. The AF models were merged with the PDB crystal structures (< 2 angstrom resolution). The merger resulted in 192 non-redundant structures. This population was filtered down to 9 structures in two steps (**Fig. 2C**). First, CD-Hit parsed the 1D sequence diversity, resulting in 59 distinct representatives. Second, DALI-based correspondence analysis resolved the 59 structures into 11 groups as assessed by a K-means clustering algorithm. These groups were screened with FunFam for functional similarity. Two groups were merged based on this screen. The final set of 9 representative structures was examined to assess whether activated CheY crystal structures, modified with phosphate analogs or activating residue substitutions, and assayed based on Y106 rotamer orientation, were segregated or randomly distributed between the 9 clusters. We found that the activated structures partitioned proportionally to cluster size, consistent with a random distribution.

The structural representatives of the 9 groups are shown in **Fig. 2D**. The representative of the dominant group is 4H6O, a *V. cholera* CheY4 protein with an isolated RR fold that does not bind the flagellar motor (Biswas et al., 2013). There are 4 other crystal structures in the set, namely 2QVG, 5TQJ, 6ONT, 5XSO. These all function as transcription factors. The remaining representatives are AF models of transcription factors. 5XSO, AF-UvrY, AF-RcsB and AF-ArcB have the RR domain embedded in a multi-domain fold. AF-NtrC is a homodimer that can oligomerize to form higher-order assemblies. These representatives reflect the dominance of transcription factors as a functional group within the RR superfamily (Gao et al., 2019).

The survey of the InterPro and PDB databases reaffirmsthat the static snapshots provided by crystal structures and AF models are not sensitive enough for a separation of active and inactive fold conformations. This result reflects the multiplicity of binding targets for diverse functions consistent with previous surveys (Gao et al., 2007). Better discrimination was obtained between the unphosphorylated and phosphorylated forms of CheY_es_ by MD simulations. The simulations supported conformational selection as a dominant feature of CheY_es_ activation (Wheatley et al., 2020).

### 3. Energy frustration in the CheY superfamily

Energy frustration provides a link for the evaluation of fold evolution. It reports free-energy scores for the native contacts relative to the decoy distribution obtain by *in silico* substitution of all amino acids for each residue position. Low residue scores indicate evolutionary stabilized contacts, whereas high scores typically reflect the energy cost for binding interactions with extrinsic peptides or small molecules. The frustration profile for the 59 post CH-Hit structures was mapped onto the individual sequences in the 1D MSA and compared against the HMM logo for sequence conservation. It was also mapped onto the structure of the major group representative (4H60) (**Fig. 3**). The map reveals (a) the RR hydrophobic core is evolutionary stable, (b) The aspartate triad (D12, D13, D57) is maximally frustrated with (c) five residues (D38, D52, G102, G105, K109) at intermediate frustration. FrustratometeR (Parra et al., 2016)reports these residue positions form water-mediated bonds, based on separation and solvent-accessible surface area (SASA). The triad participates in Mg^2+^ coordination as well as phosphate. They, together with K109, comprise 4 of the 5 conserved residue position essential for chemotaxis, D38 contributes to the solvation of the phosphorylation pocket (Hamid et al., 2022), while G102, G105 bookend the α4-β5 loop that is immobilized when target peptides associate with the binding surface (Wheatley et al., 2020). Thus, the *in silico* substitutions suggest that the energy cost for conservation of chemotactic function is maintained for RR functions. Part of the cost is due to the target binding surface, but the major contribution is due to interactions of the phosphorylation pocket with the solvent, metal ions and the phosphate as detailed below.

**Fig. 3.**
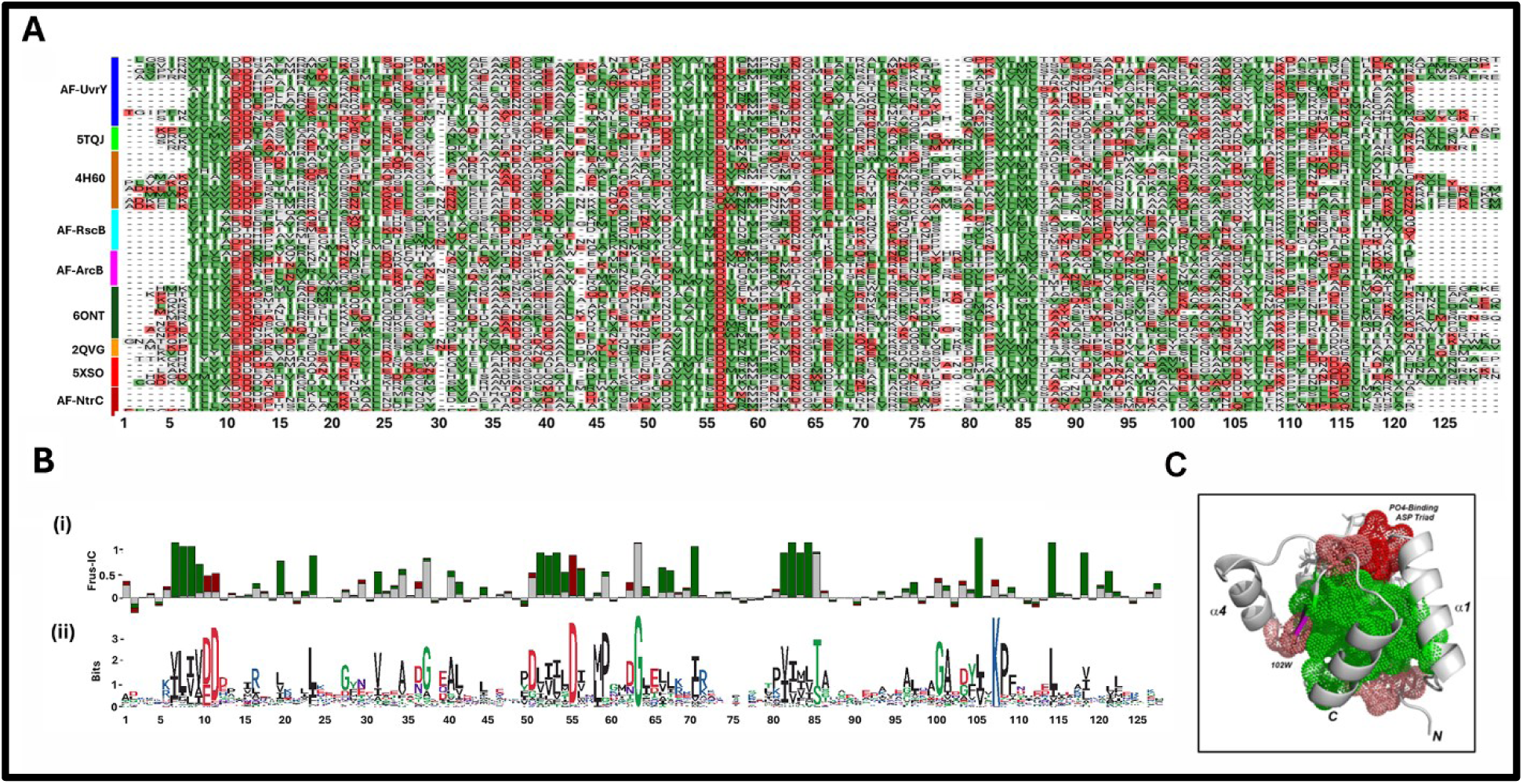
Determinants of fold stability, phosphorylation site coordination and signal transduction. **A. (i)** FrustratoEVO MSA of the 59 CD-Hit cluster representatives from the 196 member RR database colored by local energy frustration (maximal (red), minimal (green), neutral (gray)). **(ii)** The normalized maximal frustration scores as a function of MSA residue position. **B.** The histogram of normalized energy frustration scores **(i)** and the residue conservation **(ii)** as a function of MSA residue position. **C**. Frustration scores mapped onto the representative of the largest cluster (4H60 (Fig.2Cii)). Minimally frustrated (green (>0.5)). Maximally frustrated (salmon (>0.35<0.5), red (>0.5)). 4H60 N-terminus (N), C-terminus (C). Aromatic residue (W102) at target-binding interface, is equivalent to 2CHE Y106. Supplementary **Video S1** has a 3D-perspective.

### 4. Bound water at the phosphorylation pocket

The Mg^2+^ coordination dynamics in the presence and absence of aspartyl phosphate for the energy-minimized 2CHE and 1FQW_PO4_ structures are shown in **Fig. 4A**. In the basal state (2CHE), the Mg^2+^ bonds with the D12, D13, D57 triad as reported (Stock et al., 1993). Two water coordination shells were identified in the crystal structure. The 1^st^ water shell, closest to the Mg^2+,^ was reduced from three to two in 1FQW_PO4_, as the phosphate replaced one of the waters in the 1FQW crystal structure. Therewas a slight, but persistent, difference between the Mg^2+^ coordination geometry. The oxygen atom positions of the first shell water had the lowest B-factors. The three 2^nd^ shell waters had higher B-factor values comparable to the mean. The higher B-factor values in the 1FQW structure (2.4 angstrom) relative to 2CHE (1.8 angstrom) probably reflect the resolution of the electron density maps.

**Fig. 4.**
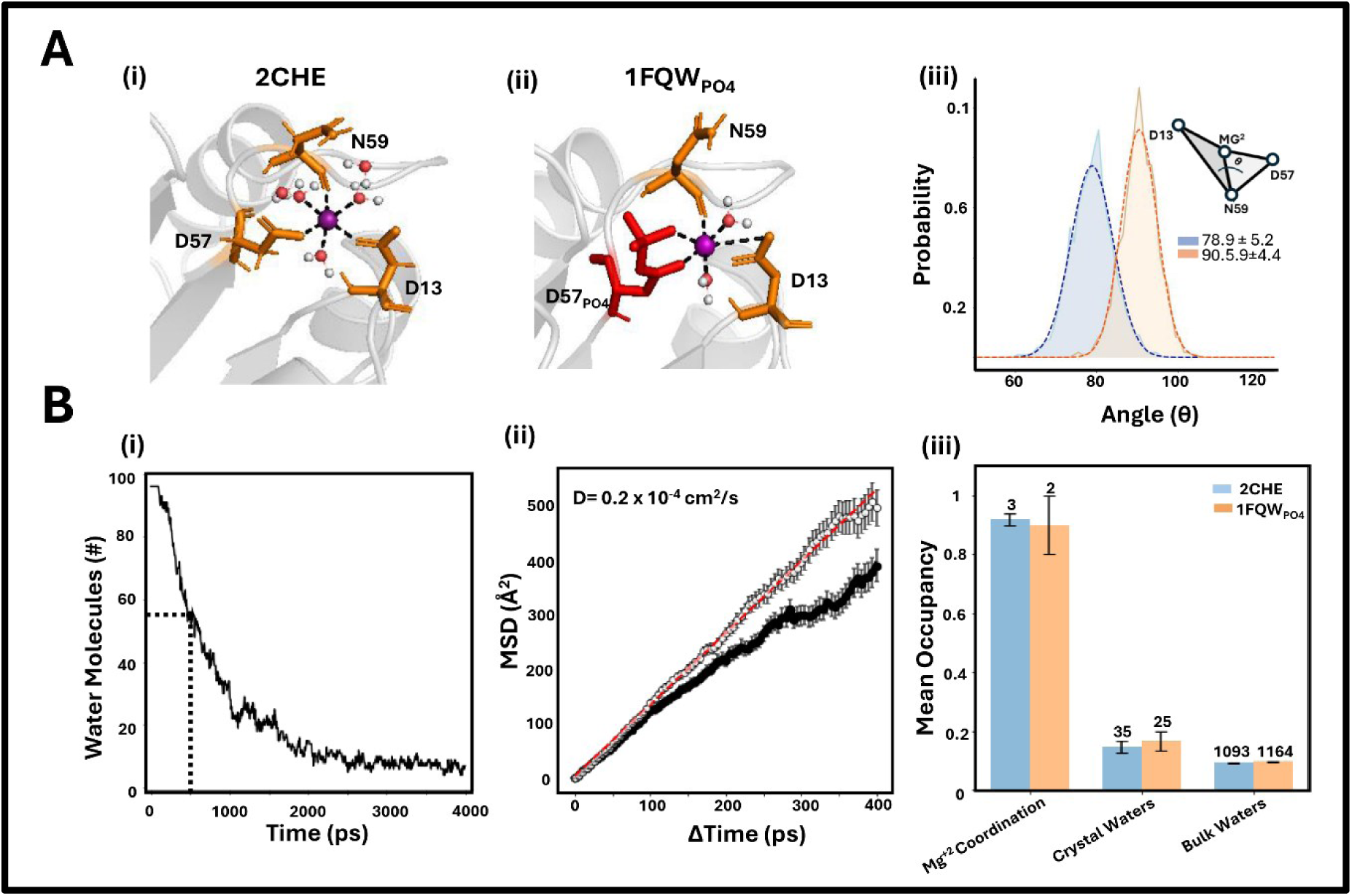
A. Mg^2+^ coordination chemistry. The **(i)** basal (2CHE) and **(ii)** phosphorylated (1FQW_PO4_) states. **(iii)** The octahedral coordination topology as assessed by the divergence angle (θ) between the planes formed by the Mg^2^ with the atomsof D13, N59 and D57,N59 is shown for 2CHE and 1FQW_PO4_. **B. Brownian mobility of crystal waters**. **(i)** The drift of the crystal waters out of the water box. Half of the population exits the box in 500 ps (dotted lines). **(ii)** Mobility of crystal waters in the presence (hindered(black circles)) and absence (unhindered (white circles)) of CheY. Linear best fit (MSD = 1.3 (Δ(Time)) (red line)) for the unhindered case (red line). D = MSD/6(1.3(Δ(Time)). The D= 0.15×10^-4^cm^2^/s for the hindered population from the corresponding best fit.. **(iii)** Occupancy of the Mg^2+^ coordination 1^st^ shell, crystal (-Mg^2+^ coordination) and non-crystal waters.. The occupancy for the 2CHE Mg^2+^ coordination 2^nd^ shell waters is <0.05 (0.01). 3 replicates, 4 ns trajectories.

The MD simulations revealed that in the absence of CheY protein, the crystal water population had a half-life of 500ps within the water box. The mobility computed from the mean squared deviation (MSD) versus Δ(time) plot gave an apparent diffusion coefficient of 2×10^-4^cm^2^/s, in line with published values (Kadaoluwa Pathirannahalage et al., 2021). In the presence of protein, the diffusion coefficient was reduced due to H-bonding with CheY residues, as assessed by Bridge2. The 1^st^ shell waters in both 2CHE and 1FQW_PO4_ were fixed for > 4ns. The crystal water population minus the Mg^2+^ coordination waters was less mobile than the non-crystal water population (**Fig. 4B**). There was no correlation between the 2CHE B-factors versus the fractional occupancy computed from the MD simulations (Pearson correlation, P_cc_ = -0.21).

We analyzed the mobility of the 2CHE crystal waters in detail. With the exception of the Mg ^2+^ coordination waters, the mobility of the crystal and non-crystal water populations was indistinguishable. The second shell waters displacement distribution showed a persistent peak around 12 angstroms. We tracked the individual trajectories to understand this effect. We found that the ring of residues that solvate the phosphorylation pocket forms a local trap that slows the escape of the second shell waters from the pocket (**Fig. S2A, B**). The residence time distributions of the protein-water (P-W) and protein-protein (P-P) H-bonds were comparable. There was no correlation between the occupancy and the residence time for P-W H-bonds, in contrast to P-P H-bonds (**Fig. S2C, D**).

The residues that form P-W H-bonds are substantially more restricted than the number of participating water molecules. As expected, charged and polar residues (n = 54) dominated this residue population (n=61), shown in the 3D contact map filtered by persistence time (> 15 ps). The preference for residue type overrode the residue SASA (**Fig. S2E**). Additional filtration by occupancy (>0.05%) highlighted the ring of acidic residues around the phosphorylation pocket.

### 5. Construction of protein-water H-bond networks

The betweenness centrality profiles of the 2CHE and 1FQW_PO4_ CheY_es_ protein-water H-bond networks, determined using the Bridge2 algorithm (Siemers and Bondar, 2021), were compared with the C^a^ centrality profiles based on the conformational entropy of torsional angles (McClendon et al., 2012) (**Fig. 5A)**. We used custom scripts to map the networks (see **Methods**). The scripts were validated by comparison against the network output by the Bridge2 GUI in the former case, and the test dataset supplied in the latter case. We classified the central Bridge2 residues into four H-bond groups to track their water bonding propensity: **(i)** residue-residue H-bonds (R-R_S_), **(ii)** residue-water H-bonds (R-W), **(iii)** switchable H-bonds that bonded alternately to multiple residues (R-R_M_) or alternately (**iv**) with water or residue (R-R/W). The (> 15 ps) persistence filter eliminated transient H-bonds that resulted from high-frequency secondary structure fluctuations and interactions of surface residues with bulk water. A second filter removed bonds with occupancy < 0.05. The C^a^ mutual information network reported modest increases in the centrality of the sequence segments around D57 (phosphorylation site) and Y106 (binding surface) in 1FQW_PO4_ relative to 2CHE. The Bridge2 H-bond networks were dominated by two sequence segments – the arginine-rich R18-R22 and the acid-rich N31-D41.

**Fig. 5.**
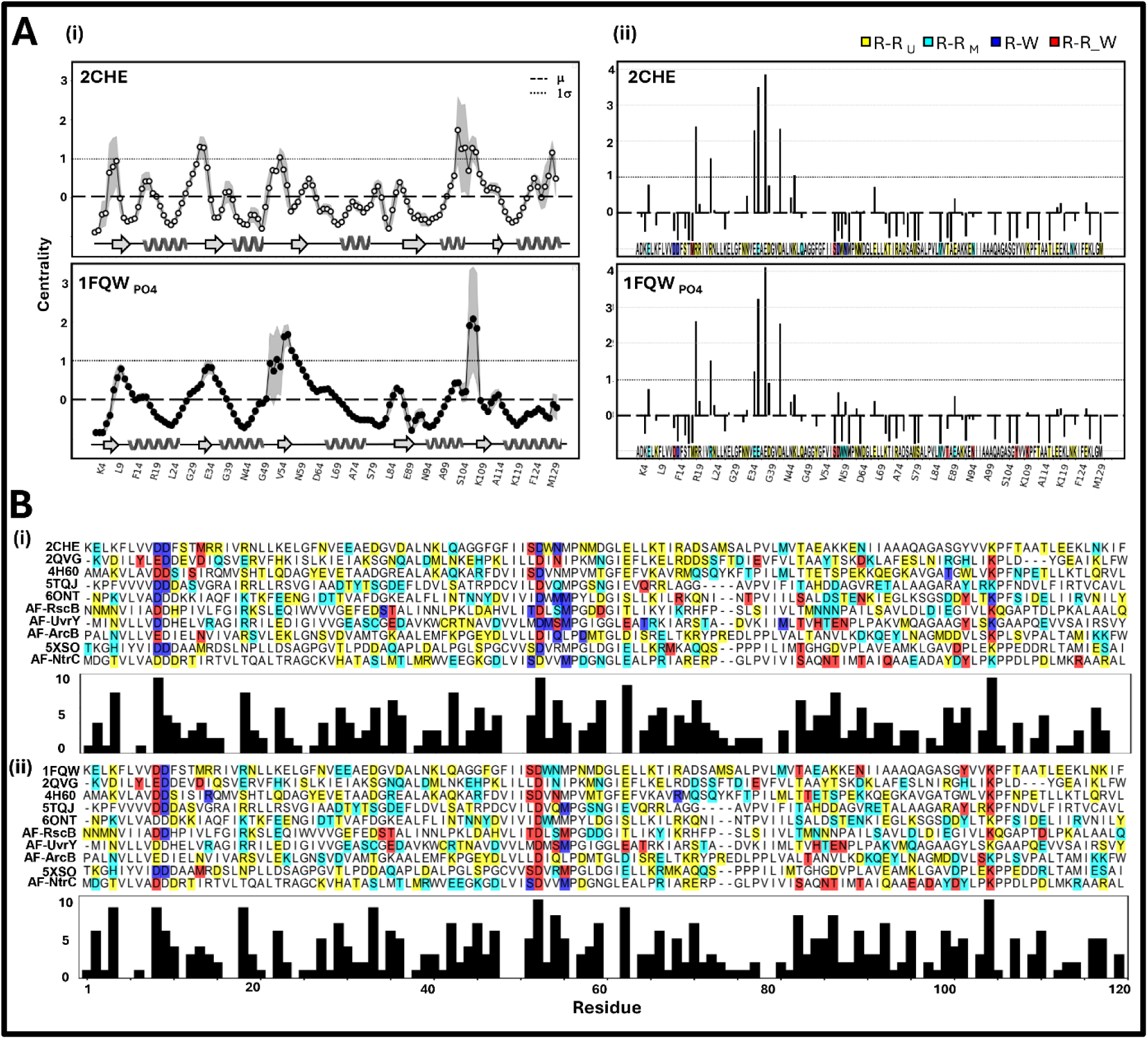
A. Differences in the **(i)** C^a^ backbone centrality and **(ii)** protein-water H-bond networks between the basal (2CHE) and phosphorylated (1FQW_PO4_) CheY states. The backbone differences were not significant as reported by one-way ANOVA (F(1.28) > F_crit_, (3.11)). Central residue positions in the H-bond network are colored by group (R-R_S_ (yellow), R-R_M_ (cyan), R-W (blue), R-R/W (red)). The R-W, R-R/W groups had higher normalized SASA (0.37) than R-R_S_,R-R_M_ (0.25). The difference was significant (p = 0.0003 (two-tailed t-test)). **B.** MSAs showing central residue position, color coded by group as in A(ii) (top) and conservation (bottom) in the H-bond networks of the RR superfamily representatives. **(i)** basal and **(ii)** phosphorylated conformations.

Bridge2 was subsequently applied to the other nine representatives of the RR superfamily ( **Fig. 5B**). In all cases, as seen for the enteric CheY proteins, the D12, D13, D57 triad binds water regardless of activation. In the CheY_es_ basal conformation, the phosphorylation pocket S56 and N59 residue positions have high water H-bond propensity. The propensity is maintained in the phosphorylated conformation for S56 but not N59. The overview of the representative RR MSA suggests that this might be a general trend modulated by residue type (S, T56, N, Q59). Mutations at both residue positions affect chemotactic performance (Pazy et al., 2009; Roman et al., 1992).

The centrality plots and MSAs overview the architecture, but do not give insight into the H-bond chemistry or the temporal coupling between residue positions. The phosphorylated D57 flips from the R-W to the R-R/W group, triggered by additional bonds formed by the phosphate, as also found for other RRs in the MSA. Representative H-bonding states in D57_PO4_ and other relay residue positions are shown in **Fig.6**. The D57_PO4_ phosphate oxygens act as both H-bond donors and acceptors. The K109 and T87 are important partners, as noted in section 1. The RR MSA shows that K109 is a water-rich residue position. The K109 ε-amino nitrogen also has high H-bond propensity. It can H-bond simultaneously to D13 and M17, as well as the D57_PO4_ phosphate. In addition, it can add an H-bond with water. T87 also H-bonds with water when bonded to D57_PO4_ in the aqueous environment of the phosphorylation pocket. Y106 rotates inwards upon phosphorylation to bond to N94. Its sidechain phenol bonds to water, possibly to reduce the energy cost for burial of this polar group in the hydrophobic core. The fluorescence quenching of W58, adjacent to D57, is a classic indicator of activation (Lukat et al., 1992). Its indole ring forms weak H-bonds in D57_PO4,_ alternatively with the D64 carboxyl oxygen and the M85 sulfur. In 2CHE, W58 only bonds with the M85.

**Fig. 6.**
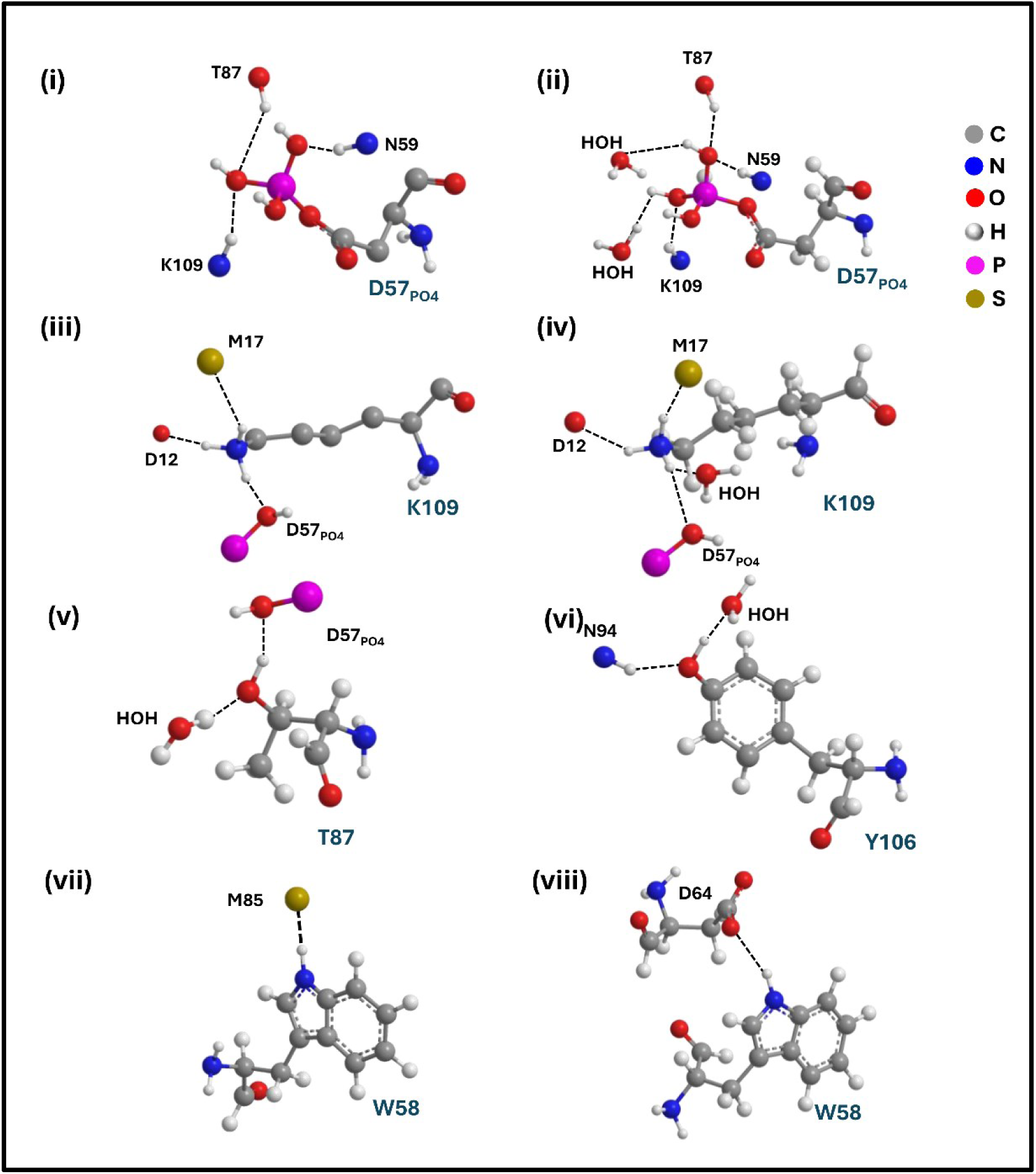
1FQWPO4 allosteric relay residue multivalent bonding states. D57PO4 (**i** (32%), **ii** (5%)). **K109** (**iii** (37%), **iv** (5%)**). T87 (v** (10%)**), Y106 (vi** (5%)). Panels (ii, iv, v, vi) show bonded waters. The mean K107-M17 persistence time in the panel (iv) state is short (< 15 ps). **W58** panels show alternate bonding states with M85 (**vii** (24%)) and D64 (**viii** (15%)). These states, except the W58-T87 state (vii (14%)), were not observed in 2CHE simulation runs. Instead, D57 bonds with water (61%), K109 with M17 (32%), and T87 with N94 (12%) in these runs. The percentile occupancies are bracketed. ChemDraw illustrations. The bond agreement between Bridge-2 and ChemDraw was >85%.

### 6. H-bond Network Dynamics

The H-bond fluctuations and their coupling with C^a^ backbone motions/sidechain rotations were effectively visualized as videos. Each video is composed of a selection of snapshots that capture all the switching transitions observed for the ((iii) R-R_M_), ((iv) R-R/W) group residues. The representative snapshots (**Fig. 7**) illustrate the distance and angle-based Bridge2 bond assignments. We partitioned the bonds as either “stable” or “switchable”. We defined switchable H-bonds as those that exchange binding partners (either residue or water) to access multiple states (n>2). Nodal residues that form single H-bonds with protein ((i) R-R_S_) or water ((ii) R-W) will fluctuate between two states – bound (ON) and unbound (OFF). Nodal residues that form multiple H-bonds ((iii) R-R_M_), ((iv) R-R/W) will fluctuate between multiple bonded states as well as the unbound state. The videos illustrate, in addition, how the fluctuations in the H-bond network are coupled to secondary structure C^a^ backbone dynamics and sidechain rotation. They provide the spatiotemporal detail into the dynamic differences between the basal and phosphorylated CheY states.

**Fig. 7.**
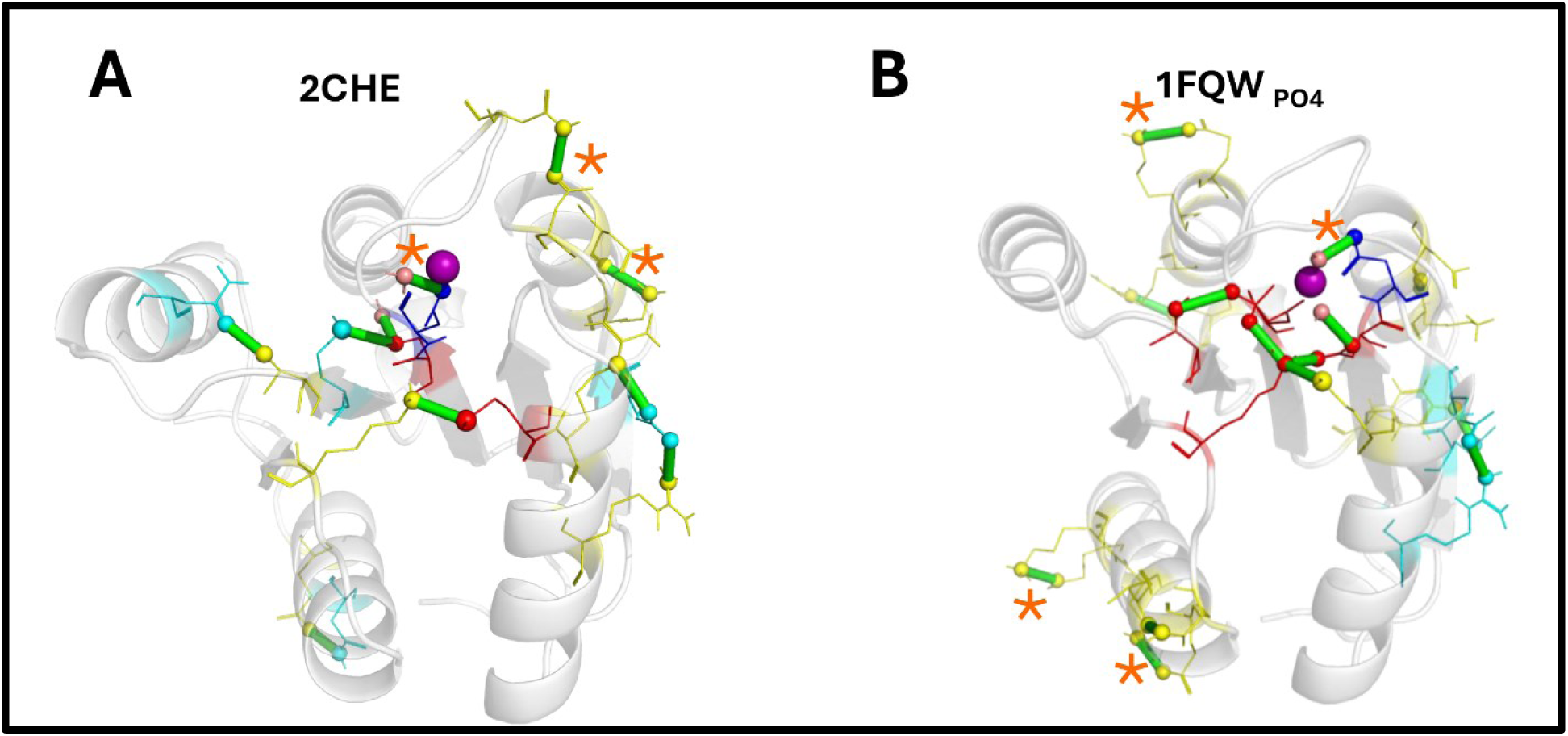
A, B: Fluctuations in the H-bond network. **A.** 2CHE. **B.** 1FQW_PO4_. Selected snapshots from videos S1, S2 (60 ns) illustrating the switchable H-bonds in residue groups ((iii) R-R_M_); ((iv) R-R/W). Sidechains (line representations) of the Bridge2 central residues with the bonding atoms (spheres) color-coded by group as in Fig. 5Aii. Water (salmon). H-bonds (green lines). Asterisks mark stable bonds between residue groups ((i) R-W; (iii) R-R)

### 7. Comparison of the physical H-bond networks

The 2CHE, 1FQW_PO4_ protein water networks are compared in **Fig. 8**. The largest network community in both cases is water-mediated with conserved residues in water groups (ii) and (iv) including the D12, D13, D57 triad localizing to the phosphorylation pocket {(2CHE (D12, D13, D57, N59, W56, W58, M85, M17, K109)), (1FQW_PO4_ (D12, D13, S56, D57, W58, N59, D64, M85, E89, K91, N94, Y106, K109). However, the residue water H-bond propensity and the coupling between them, as measured by the occupancy, change. In 1FQW_PO4_, the centrality of the common residues remains relatively unchanged, but the community expands to include more residues, and the occupancy of the bonds between D12, D13, D57, T87, and K109 increases to dominate the network.

**Figure 8.**
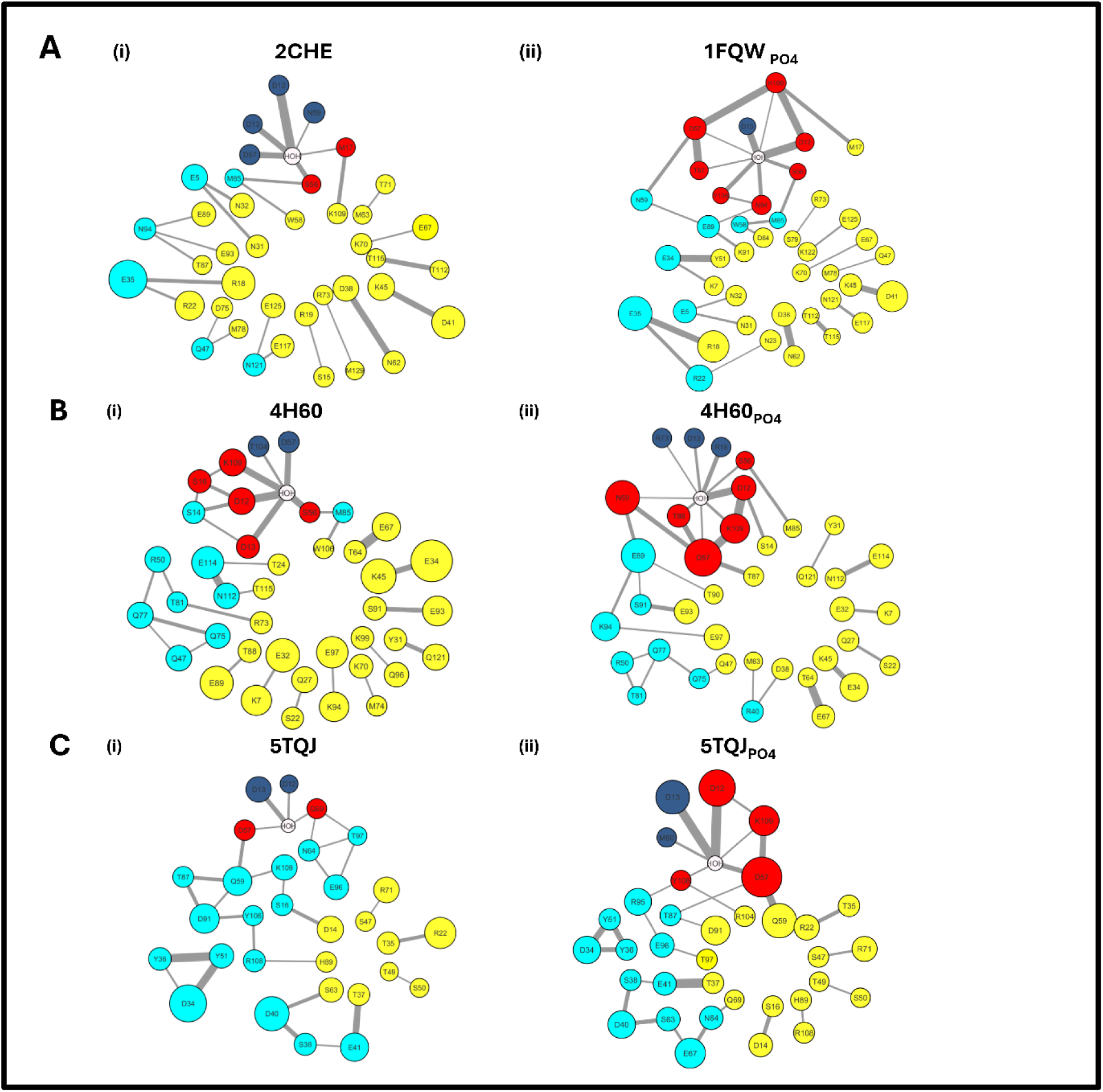
Bridge2 networks of representative response regulators. **(A)** CheY_es_ **(i)** basal (2CHE), **(ii)** phosphorylated (1FQW_PO4_). **(B)** 4H60 **(i)** basal**. (ii)** phosphorylated **(C)** 5TQJ **(i)** basal **(ii**) phosphorylated. Residue nodes are colored by group, as in Fig. 5Aii. with size indicating centrality, while the thickness of the edges between them reflects bond occupancy Cumulative 3×20ns replicates.

We extended the analysis of the enteric CheY templates to the basal and phosphorylated states of the representatives of the two major RR sub-families (Fig. 2Cii), namely 4H6O and 5TQJ. 4H60 (*V. cholerae* CheY4) represents the largest sub-family that includes structures of chemically modified phosphate mimics as well as intraspecies paralogs such as *V. cholerae* CheY3 that binds FliM, in contrast to CheY4. Other members include enzymes such as CheB and NtrC, and transcription factors (<10%). 5TQJ is a *Burkholderia phymatum* transcriptional regulator that belongs to the LuxR RR sub-family (Tsai and Winans, 2010). It may dimerize.

### 8. The coupling between residue H-bond states

The physically-connected H-bond network will have associated with it a dynamically coupled network that reflects the backbone and sidechain dynamics. The dynamic network, unconstrained by distance includes significant long-range (12 angstrom (= 3.5 x H-bond) mean distance) couplings. We measured the mutual information (nMI) for these dynamic couplings between residue H-bond states, after removal of the H-bond contacts, if any, between the paired residues (R1, R2) (see **Methods**).

The significant couplings (> mean + σ of the non-zero MI distribution) for the basal and phosphorylated CheY and *V. cholerae* CheY4 are shown for the Bridge2 central residue matrices (**Fig. 9A**). A notable difference between the basal (2CHE) and phosphorylated CheY (1FQW_PO4_) was the increased couplings involving T87 and Y106. For *V. cholerae* CheY4, phosphorylation increased the strength and number of couplings involving T87 and K109. Couplings involving W102 (= CheY Y106) were not significant for either the basal or phosphorylated conformations.

**Fig. 9.**
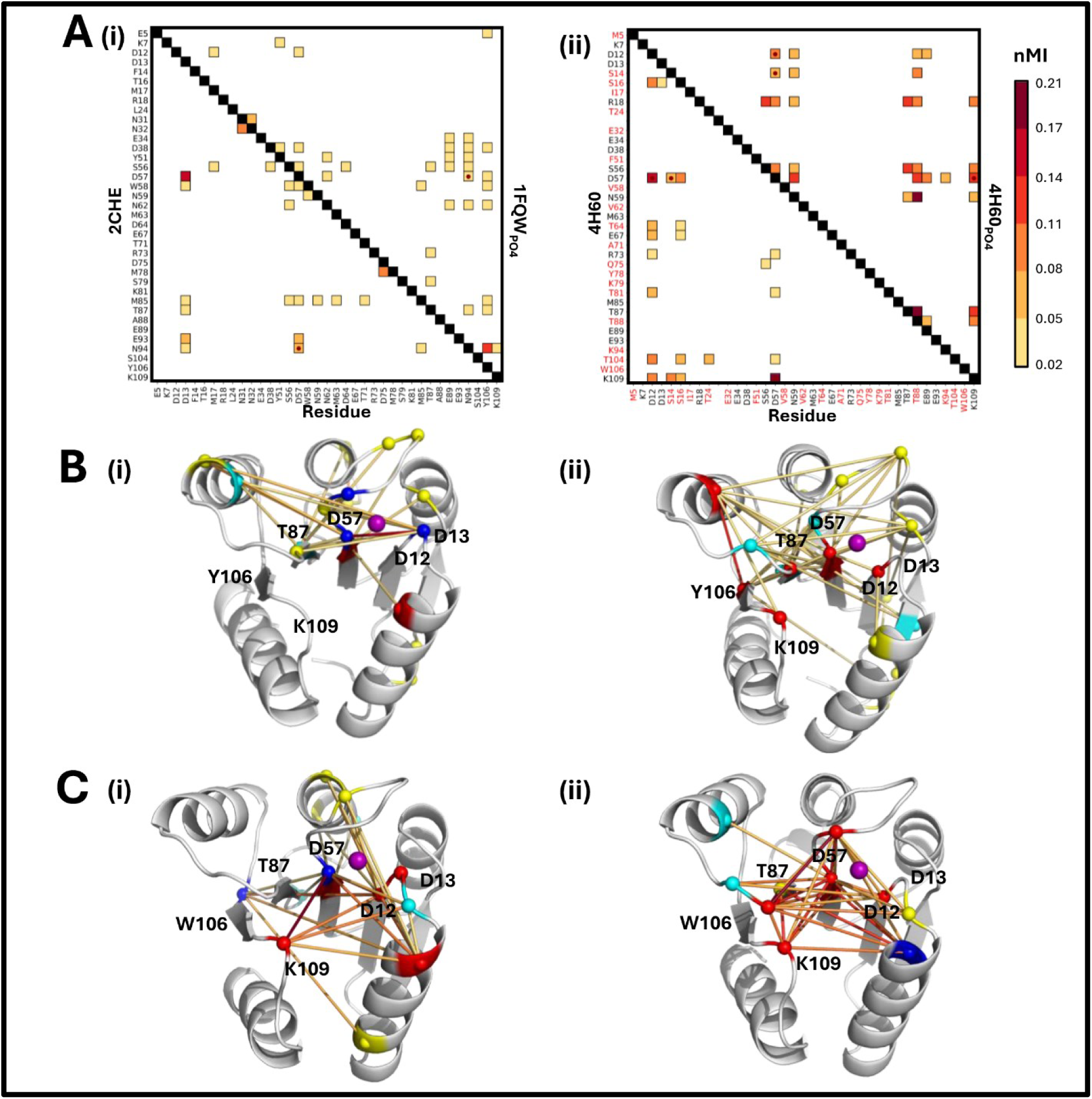
The coupling between chemical group state transitions. A. Difference maps between basal and phosphorylatedforms.(i) S. Typhimurium apo(2CHE) conformation/ E. coli phosphorylatedconformation (1FQW_PO4_). **(ii)** V. Cholerae basal / phosphorylated CheY4. Residue (red font) mark positions with residue substitutions relative to S. Typhimurium CheY. Fill color indicates nMI value (Bar). **B,C**. The significant nMI couplings mapped onto the 3D-structures of – B.**(i)** basal (10.4±1.1), **(ii)** phosphorylated (13.9±1.2) CheY_es_. **C.(i)** basal (14.2±1) **(ii)** phosphorylated (9.5±0.6) V. cholerae CheY4. Numbers in brackets denote C^α^-C^α^ distances, in angstroms, of the coupled pairs. Line color scales with nMI value, as in A. Residues are colored by group as in Fig. 5Aii.

**Fig. 10.**
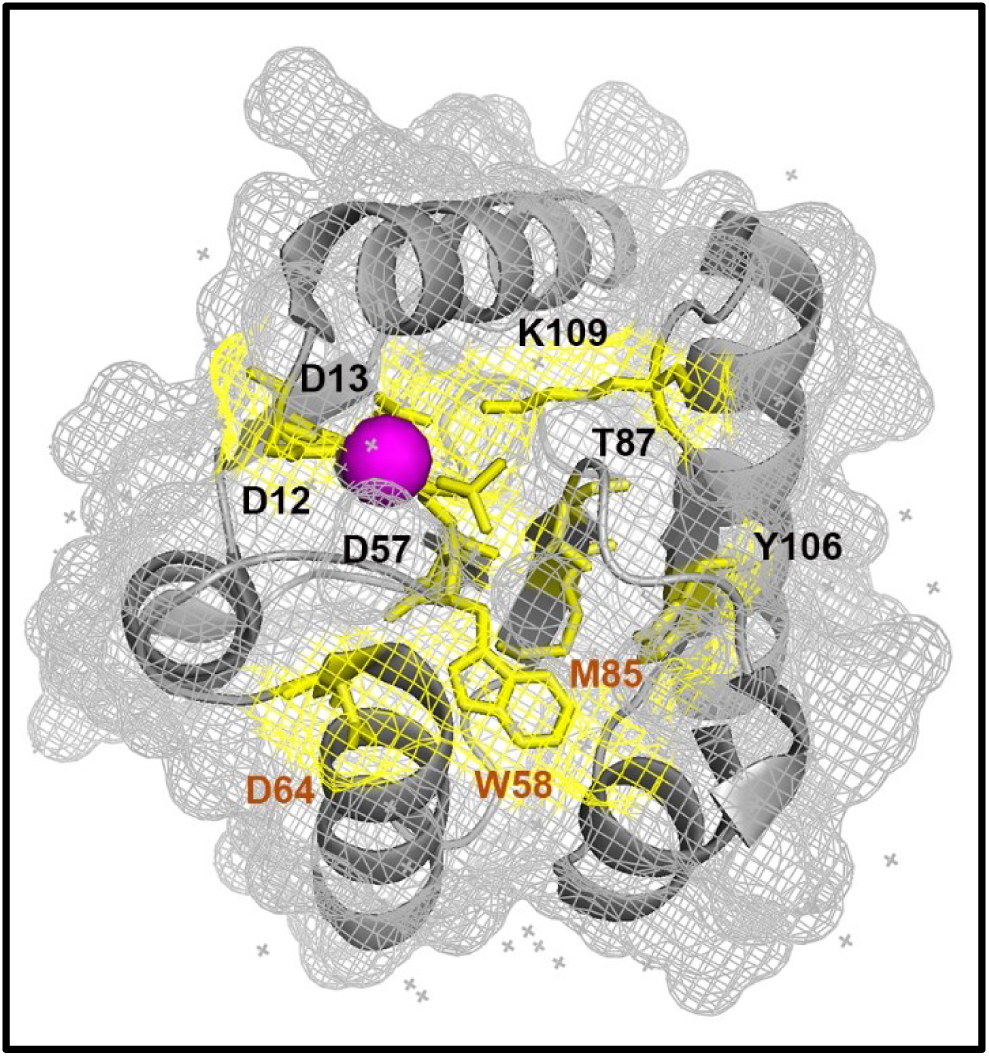
H-bonds in the water mediated CheY_es_ relay. The 1FQW crystal structure has underpinned the knowledge of CheY_es_ activation. Therole of H-bonds, including P-Wbonds, revealedby this studyis mapped (yellow mesh) onto the static structure.Mg^2+^ (magenta sphere). Water mediated phosphorylation triad and relay (black labels). W58 bridge (orange labels).

**Fig. 11.**
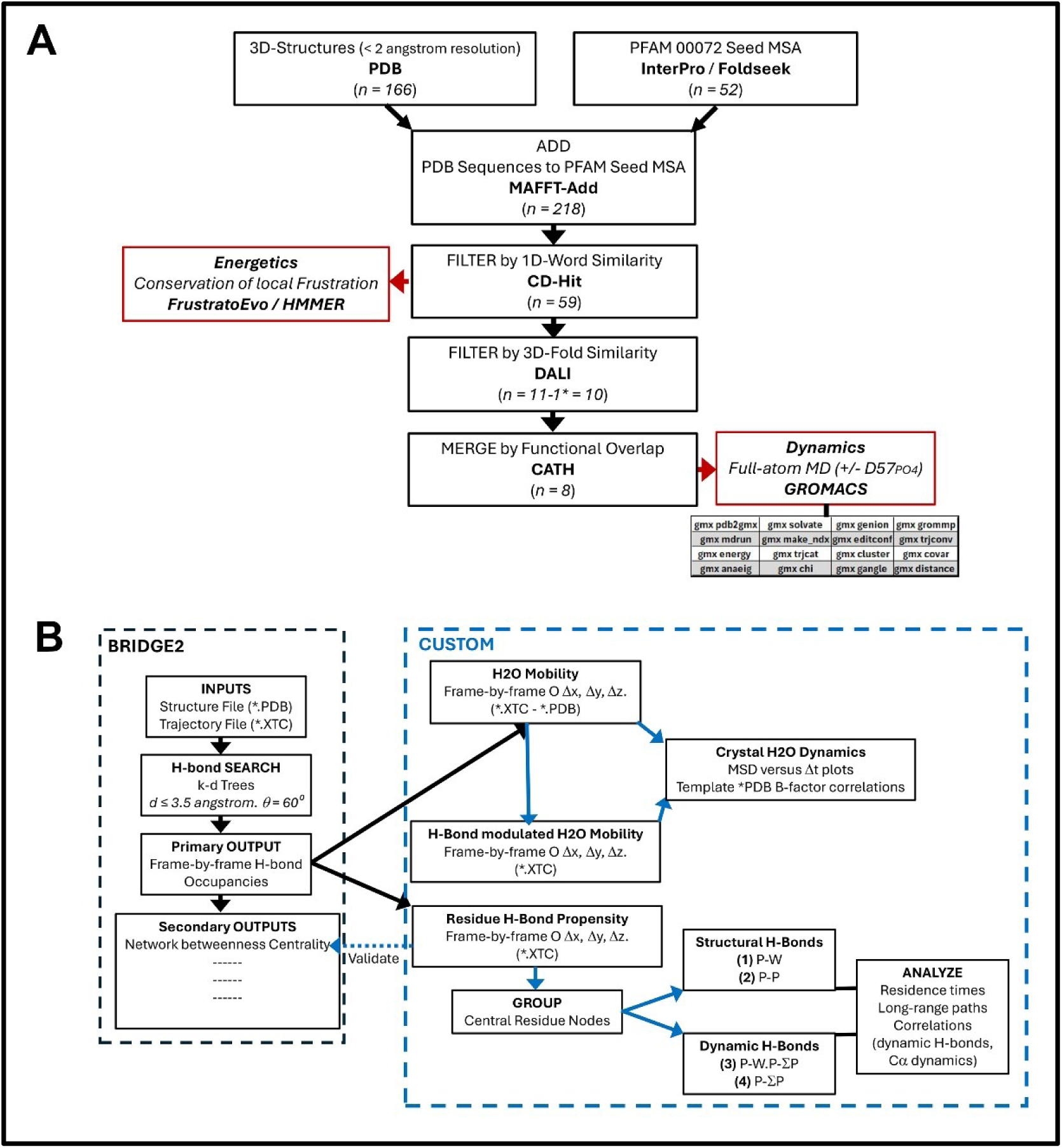
A. RR superfamily construction filtration and analysis. The workflow for the construction of the composite 1D sequence and associated 3D structure database and the 4 stages for selection of structures for analysis is shown. For simulations of the dynamics, each structure selection was simulated with and without the phosphoaspartyl modification in the presence of Mg ^2+^. See text for the GROMACS operators used for the MD simulations and analysis. **B. The Bridge2 H-bond protein-water network.** The workflow for the metrics extracted from the Bridge2 output.

Residues can form H-bonds with multiple partners (see Fig. 6). We will use the acronym “*H-bond_MAX_*“ for the maximum number of H-bonds formed by a residue. The number of bonding states per residue will increase with *H-bond_MAX_* combinatorially,given that residues can simultaneously form H-bonds with multiple partners. Another consequence of phosphorylation, in both CheY and *V. cholerae* CheY4, was the increase in the number of states. D57_PO4_ had a greater *H-bond_MAX_* than D57, resulting in a marked increase in the state multiplicity. The K109 *H-bond_MAX_* also increased upon phosphorylation due to the compaction of the protein core (**Fig. S3**).

The phosphorylation pocket forms the central community node for the basal and phosphorylated conformations in both proteins, as appreciated by the map of the dynamics network onto the 3D structures. In both cases, the community number and strength increase with phosphorylation. (**Fig. 9B, C**).

## Discussion

The CheY_es_ core has an extensive H-bond network with significant contributions from bound water molecules (Volz and Matsumura, 1991). Differences between H-bond interactions have been visualized in inactive versus active crystal states. CheY and other RRs are likely to exist in their Mg^2+^ bound form *in vivo* since Mg^2+^ is the most abundant divalent cation in prokaryotes (Franken et al., 2022). Mg^2+^ coordination changes the bonding between the phosphorylated D57 site and adjacent conserved residues (D12, D13, K109) (Stock et al., 1993). BeF_3_ elicits further changes in residue and water interactions around D57 (Cho et al., 2000; Lee et al., 2001). MD simulations have described the CheY_es_ C^α^ conformational landscape guided by an extensive CheY_es_ database of mutant crystal structures, with *in silico* phosphorylation used to compare CheY_es_ with other RRs in one study (Foster and West, 2017).

These seminal insights into the activation mechanism are nevertheless incomplete. Molecular CheY_es_ structures have been correlated with behavioral phenotypes, but there is a sharp decrease in such knowledge for other RRs. Importantly, water dynamics are required even in CheY_es_ for the characterization of the phosphorylation pocket and the allosteric H-bond network linking it with the target binding surface. The unambiguous determination of H-bonds and water molecule positions in experimental structures is beset with challenges (Prisant et al., 2020; Schiebel et al., 2018), in spite of notable successes (Giannetti et al., 2025; Tani and Fujiyoshi, 2014).

Here, we used MD simulations, supplemented with automated, high-throughput H-bond detection (Siemers and Bondar, 2021), to study the CheY_es_ basal and phosphorylated conformations. We exploited the FrustratoEvo algorithm to reveal energy cost for the conservation of the phosphorylation pocket and the target binding surface across RR superfamily members. We segmented the H-bond networks into groups to parse plastic from hard-wired H-bonds to reveal generic and plastic elements of the molecular network across different functions by comparison of two RRs with different functions from CheY_es_. Finally, we constructed coupled dynamic networks to evaluate the protein response to unitary H-bond transitions. Our methodology is not without its limitations, as noted in Methods, but it provides a novel perspective for the study of RR activation.

### The stable core

The hydrophobic core of the RR superfamily was defined by its minimal energy frustration profile (Fig. 3). It offers one route for connecting the phosphorylation pocket to the target binding surface that may regulate the phosphorylation kinetics of chemotaxis response (Foster et al., 2021; Hamid et al., 2025) and perhaps other RRs. Our focus here has been the H-bond network. Stable elements of this network predominantly form intra-helical hydrogen bonds between helices α1, α2, and α5, largely insensitive to the transition between the basal and phosphorylated conformations.

### The phosphorylation pocket

The phosphorylation pocket consists of the aspartate triad (D12, D13, D57), residues involved in metal cofactor coordination (e.g. N59) and solvent. It is the major community node in CheY. Phosphorylation triggers reorganization of its P-W H-bond network. These changes have been reported in published crystal structures, but the present study adds temporal detail to assess the coupling between key bond transitions, characterizes the role of the solvent in Mg ^2+^ coordination and reveals how both Mg^2+^ and the PO_4_ strengthen the pocket to orchestrate allosteric communication with the target interface. The conserved D12, D13, D57 triad is “water-rich” (>50% membership in P-W H-bond groups ii, iv) with maximal energy frustration across the superfamily. A chain of H-bonds connects D57 with the interfacial Y106 via N94, E89, and N59. N59 was one of the residues implicated in the steric block of D57 from solvent attack (Pazy et al., 2009). It has high solvent accessibility as reported by XFMS experiments (Hamid et al., 2022). Threonine H-bonds tighten the association of secondary structure elements in glycophorin A (Smith et al., 2002), and T87 could play a similar role in CheY_es_. The fluorescence quenching of W58 is a classic indicator of activation. We find that the W58 indole group can bridge between β4 (D64) and β5 (M38) when reoriented by phosphorylation. Tryptophan indole H-bonds are weak in a high dielectric, but the internalized W58 may participate in communication via the hydrophobic core to the target interface, as documented for membrane proteins (Khemaissa et al., 2022).

### The allosteric relay

Allosteric communication between the phosphorylation pocket and the target binding surface is mediated by T87 and K109. In the basal CheY state, T87 is part of a small local network, while K109 hydrogen bonds with M17 at the edge of the phosphorylation pocket. Upon phosphorylation, it binds D12 and D57 to form an expanded phosphorylation pocket together with T87. Both residues H-bond with water. K109 can achieve an *H-bond_MAX_* of 5 H-bonds as reported for lysine with backbone carbonyl atoms (Rogacheva et al., 2017). Thus, the expanded phosphorylation pocket community network is enriched in structural waters as well as in coupling strength.

Y106 at the target binding surface is exposed to solvent in the basal state and is not central to the protein water network. It internalizes to transition to a central residue in the network upon phosphorylation. It’s hydroxyl group H-bonds with solvent in this conformation. This bond could reduce the energy cost for burial of the hydroxyl in the hydrophobic core (see (Pace et al., 2001)). Importantly, there is a dynamically coupled network that is responsive to fluctuations in the H-bond network due to bond associations / dissociations. Y106 communicates with the phosphorylated pocket via multiple residues (N94, T87, M85, N62, D38) as part of the dynamic network.

K91 acetylation mimics D57 phosphorylation (Fraiberg et al., 2015). and immobilize the β4-α4 loop in the 13DK106YW phosphomimic when bonded consistent with reduced solvent accessibility (Wheatley et al., 2020). It is not part of the basal H-bond network but connects to both D57 and Y106 in the phosphorylated network via N94, E89, and N59. N59 was one of the residues implicated in the steric block of D57 from solvent attack (Pazy et al., 2009). It has high solvent accessibility as reported by XFMS experiments (Hamid et al., 2022). Key elements of the water-mediated relay are mapped onto the 1FQW structure. Their contiguity depends on protein motions, ranging from single H-bond association/dissociations to a-helix packing interactions whose appreciations require atomistic simulation of the dynamics.

### Functional adaptation

A number of bacterial species have multiple CheY homologs involved in chemotaxis. *V. Cholerae* provides one example. *V. cholerae* CheY3 binds the flagellar motor protein FliM, but *V. cholerae* CheY4 doesnot. It is thought that the latter serves as a phosphate sink to optimize the timing of the chemotactic response (Biswas et al., 2013). The equivalent residue position to CheY Y106 in *V. cholerae* CheY4 is W102. Study of its structure (4H6O.pdb) reveals that W102 does not contribute to the H-bond or the associated dynamically coupled network in either the basal or phosphorylated conformations, consistent with the absence of interaction with the FliM motor protein. Nevertheless, the phosphorylation dependence is conserved. The phosphorylation pocket is the central community in the H-bonded networks. In the basal conformation it comprises D12, D13, S14, S16, K109. It expands upon phosphorylation to include N59, T87, T88, with D57 as themajorresidue node, connected to N59, T87 and K109. N59 connects to E89 that is the organizing node for a smaller community network. In the associated dynamically coupled network the T87 – E89 segment transitions to a major node upon phosphorylation, with an increase in K109 connectivity.

We also studied the *Burkholderia phymatum* transcriptional regulator structure 5STJ. The 5STJ phosphorylation pocket is the central community analogous to the CheY and *V. cholerae* CheY4 H-bond networks. Interestingly, in the basal conformation, D57 is connected via Q59 to T87, D91, Y106 and K109 at the binding surface. Upon phosphorylation, D57 connects directly to Q59, T87 and K109. The D57 – Q59 and D57-K109 couplings dominate the network. Thus, the central features are conserved in the 5STJ network, even though dimerization, if the functional response, may be slower than chemotactic excitation. The anomalous D57 connectivity in the basal conformation may link to response timing, but validation will require experimental data.

### Implications of the study

An understanding of water dynamics has facilitated drug design as documented by a growing number of studies (Samways et al., 2021; Spyrakis et al., 2017). Drug development against RRs is an attractive proposition since this superfamily does not have mammalian homologs. It has been difficult since available metrices have had limited success in discrimination between subfamilies. It should be facilitated by the tools developed in this study that, as far as we are aware, is the first comprehensive examination of protein-water H-bond dynamics in bacterial RRs.

## Materials & Methods

### Databases & Webservers

Sequences and structures were downloaded from (InterPro, UniProt) and (PDB. Foldseek) respectively. The PFAM PF00072 RR seed MSA was expanded with the addition of the downloaded structures sequences. Online servers for additional MSA and manual curation plus 3D-structure alignment, modification and in-silico mutational analysis of single residue positions are listed in **Table 1 (Webservers / Databases)**. Software installed in-house for sequence analysis, structural dynamic analysis and visualization, statistics and figure preparation is listed under **Table 1 (Tools).**

**Table 1.**
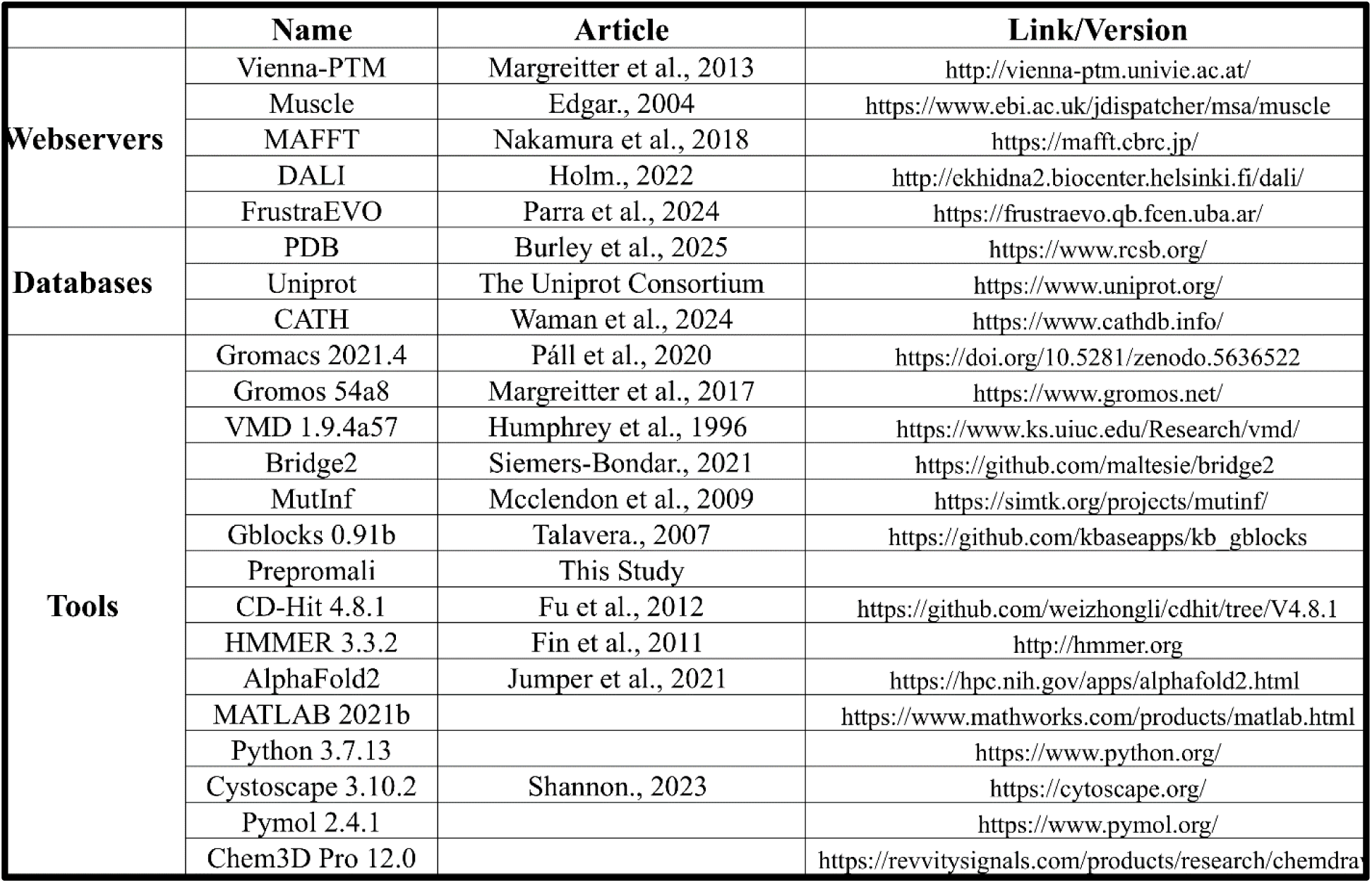

### Structural Variation in the RR superfamily

The MSA of the Pfam PF00072 response regulator receiver domain seed sequences (n=52) was used as the template. The sequences for RR crystal structures from the Protein Data Bank (PDB) with better than 2.0 angstrom resolutionwere added to the PF00072 seed MSA with the MAFFT-Add utility (Nakamura et al., 2018). The crystal structure database was a heterogenous mix of metal (Mg^2+^, Mn^2+^) bound and free basal and activated state structures. Activated structures were defined as those that contained BeF_3_ or D57 modified activation mimics. Foldseek AlphaFold models (Jumper et al., 2021) of the PF00072 sequences were combined with the PDB structures to create a 3D structure dataset to match the combined MSA. The MSA was preprocessed by trimming to the template CheY sequence with a custom code (Pandini et al., 2015), here termed PreProMALI (Preprocessor for Multiple Alignment), to eliminate residue positions absent from the PDB query sequence, 2CHE.pdb, the basal state template for the RR domain. The trimmed sequences were clustered based on 1D-sequence similarity. We selected CD-Hit (Fu et al., 2012), among several options (Pavlopoulos et al., 2023), based on previous experience with it on datasets of similar size (Khan, 2022). CD-Hit is based on the principle of short word filtering that circumvents assumptions used for sequence alignment algorithms to identify and cluster similar sequences (Li and Godzik, 2006). The cluster representatives were analyzed for local frustration with FrustratoEvo (Freiberger et al., 2023; Parra et al., 2024) and filtered by fold similarity with DALI (Holm, 2022).

The FrustratoEvo server was built on the Frustratometer algorithm (Ferreiro et al., 2007; Parra et al., 2016) described previously (Hamid et al., 2022). FrustratoEvo extends the central concept of local frustration as assessed by a decoy distribution of *in silico* mutants to map the evolution of the local frustration at the single residue level in protein families. The single residue frustration information content,

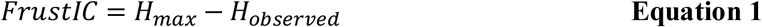

The observed conformational entropy, *H*_*observed*_, for this residue position in the MSA of family members will be

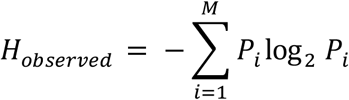

where *P*_*i*_ is the probability for the *i*^th^ (*M* =3) state (minimal, neutral, maximal) defined relative to the decoy distribution obtained by substitution of all residue types in this position with the contact and solvent accessibility values of the surrounding residues fixed (Freiberger et al., 2023). The maximum entropy, *H*_*Max*_ was computed using the non-uniform background probabilities determined earlier (Ferreiro et al., 2007), with 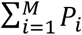 normalized to 1.

The DALI server uses intramolecular distance matrices to compare 3D-structures (Holm and Sander, 1993; Holm and Sander, 1995). It can detect remote homologs, assisted with hidden Markov models (Karplus et al., 1997), in experimental and AlphaFold predicted protein databases (Holm et al., 2023). The correspondence between structures were reported as eigenvector heatmaps and correspondence plots. The 59 CD-Hit cluster representatives were resolved into 10 groups and 1 outlier in the PC1-PC2 space using the K-means module available in the Python *Scikit-Learn (sklearn)* machine learning library. The outlier was removed.

K-means partitions the *N* structures represented by 2D coordinates in DALI eigenvector (*eigen*1, *eigen*2) space. The optimal number of clusters, *gC*_opt_ is obtained by minimization of the cost function *J*(g*C*), the aggregate intra-cluster sum of squares.

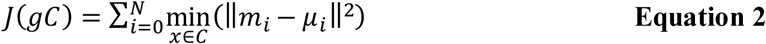

where *μ*_*i*_ is the (*eigen*1, *eigen*2) centroid of the grouped cluster, *gC*_*i*_, and *m*_*i*_ is the centroid of cluster *C*_*i*_ within *gC*_*i*_. Biological functions of the 10 representative structures output from DALI were compared with the CATH database (Sillitoe et al., 2021). The central CATH module performs structural classification of determined and predicted protein domains into homologous superfamilies (Bordin et al., 2024; Bordin et al., 2023). The superfamilies are further classified into functional families (“FunFams”) (Das et al., 2015; Rojano et al., 2022) based, in part, on the gene ontology database (Attrill et al., 2019). The FunFam functional predictions for the 10 DALI representatives aligned with function as established by experiments (Foster et al., 2021), but there was redundancy with any one sequence assigned to multiple FunFam families. This result reflected the fact that the RR fold is not a sensitive readout of its function (Gao et al., 2007). We merged two pairs of the DALI representatives with very similar FunFam profiles. Principal component analysis (PCA) PCI-PC2 plots (*gmx anaeig*) of the component members identified the average C^α^ backbone fold for each merged group. In one case, a member that overlapped with the average fold ((0,0) coordinate in PC1-PC2 space was selected as representative of one merged set. In the other case, an AlphaFold model of the average C^α^ fold was constructed by assigning the predominant residue in the group MSAto residue positions. The final 8 structures were used for molecular dynamics (MD) simulations and network analysis. The selection of the combined RR database down to the final representative structures, and the two-stage analysis of its energetics and dynamics are schematized (**Fig. 8A**). The analyses of the dynamics are detailed below.

### Molecular Dynamics

*Computer Operating Systems:* The long (1μs) simulations of the 2CHE and 1FQWPO4 template structures run on the Biowulf supercomputer with GROMACS 2024-gcc11.3 were allocated 2 compute nodes (4xNVIDIA, A100 GPUS (80 GB VRAM, 6912 cores, 432 tensor cores)). The long simulations of the 3CHY template (0.74±0.13 μs) and AF-ArcB (20 ns) were done on the NUST supercomputing facility (2 x Intel 6-core Xeon E5-2630 V2 2.60GHz, 12 cores, 64GB RAM, with 4 Nvidia-Tesla K10-GPUoption). The remaining short (20ns, 40ns) simulations were performed within a GROMACS 2021.4 installation on Ubuntu 22.04.3 on LUMS BIRL servers (4 Intel Xeon E7-8880 v3 4×18-cores, 1024GB RAM).

*Parameterization.* The BeF_3_ was removed and the D57 residue modified to dianionic D57 phosphoaspartate in the 1FQW template structure using the Vienna PTM server (Margreitter et al., 2017) to obtain phosphorylated CheY (1FQW_PO4_). The database representative structures were modified by the addition of Mg^2+^ to create the Mg^2+^-bound basal state, and modified to D57 phosphoaspartate, with the same protocol as used for the template (1FQW_PO4_), to create Mg^2+^-bound activated states. The GROMACS operations for the simulations and the subsequent analysis of the network dynamics are outlined below.

Water was parameterizedafter file conversion (*gmxpdb2gmx*) as the SPC 3-point model. The model has been studied as a function of temperature and long-range interaction cutoff (Mark and Nilsson, 2002). SPC is one of multiple water models, none of which completely reproduce the properties of water (Kadaoluwa Pathirannahalage et al., 2021). The proteins were solvated in a cubic water box with 1 nm separation between the protein and box boundaries (*gmx solvate*) and The box was adjusted to neutral pH (7.0), adjusted to 150 mM ionic strength by addition of Na^+^ and Cl^-^ ions (*gmx genion*) and checked for molar water density (*gmx energy*).

*Preproduction.* The system was equilibrated as instructed in the GROMACS tutorial (https://tutorials.gromacs.org/docs/md-intro-tutorial.html). In brief, energy minimization using the steepest descent algorithm was followed by NPT (isothermal-isobaric ensemble) (100ps) and NVT (canonical ensemble) equilibration (100 ps). Temperature control was set at 300K and 0.1 ps coupling constant with the Berendsen algorithm. Isotropic pressure was set at 1 bar with the Parrinello-Rahman barostat. Nonbonded interactions had a 1.0 nm Verlet cutoff, with the long-range interactions approximated by the particle-mesh Ewald (PME) method. We checked that the frame-by-frame root C^α^ root-mean-square deviations (RMSDs) reached a stationary value. This value, averaged over all energy minimization runs was 0.027 ± 0.003 nm).

*Production.* The production runs (*gmx mdrun*) used structure files created (*gmx grompp*) from the same parameter and topology files as the preproduction runs to process the output from the pressure equilibration. The 1μs simulations used a 2fs integration step. The 20ns simulations used an 1fs integration step to capture water dynamics. Simulations were preformed in triplicate for each structural state (3 templates plus 8×2 database representatives). Both 1μs and 20ns simulations were performed for the template structures (2CHE, 1FQW_PO4_) to assess the error due to finite sample size. The proteins were centered, then rotationally and translationally-aligned in the structure (*gmx editconf*) and trajectory (*gmx trjconv*) files. Index files (*gmx make_ndx*) edited these output files for concatenation (*gmx trjcat*), removal of water or other components as needed.

### Construction of the protein-water H-bond network

A complete identification of H-bonds and water positions based on visual inspection of protein structures is challenging so automated methods have been developed for this purpose. Here, we extended the use of Bridge2 (Siemers and Bondar, 2021) introduced in our initial study (Hamid et al., 2022) to analyze MD trajectories (**Fig. 8B**). The use of k-d trees is central to the computational power of Bridge2 that renders it two orders of magnitude faster than other options (Siemers et al., 2019) such as GROMACS *gmx hbond*. In common with other programs Bridge2 uses ≤3.5 Å bond distance and 60° angle to search for H-bonds. The frame-by-frame occupancy of the identified H-bonds is the primary output of the search. The algorithm also computes secondary metrics from this primary output -network measures, long-range contiguity, water-wires and H-bond motifs. However, the display of these metrics by the Bridge2 GUI is expensive in computer memory for large trajectory files. Therefore, we downloaded the raw occupancy data and wrote our own code to analyze and visualize metrics of interest. We validated our code by the match of the network centrality with Bridge2 using structure files alone.

We then used the Bridge2 occupancy data to, first, characterize the water-mediated coordination of the Mg^2+^ and its modulation by D57_PO4_. We computed water diffusion coefficients in the absence of protein (free mobility) as another benchmark of the SPC water model, then compared the position dependent mobilities of the waters at the Mg^2+^.D57_PO4_ binding pocket and other locations reported in the 2CHE and 1FQW crystals against the mobility of free water as well as the waters added to solvate the protein in the simulation box.

Second, we determined the protein-water H-bond network. In protein network models, the residues form nodes and the interactions between them edges. The readout of the protein-water H-bond network architecture by the Bridge2 algorithm is the betweenness centrality (Siemers and Bondar, 2021). Importantly, the Bridge2 networksare physical networks wherethe edges are set by steric constraints .( ≤3.5 Å, 60°). The residues node centrality reflects the mean occupancies and residence times of their H-bonds formed with water and other residues.

### The Analysis of Dynamic Networks

For dynamic networksthat are not subject to strict steric constraints, the correlations weremeasured as the mutual information between dynamic couplings – between residue sidechain rotations (*gmx gangle*), bond distances (*gmx distance*), backbone dynamics (*gmx chi*). The master equation for the normalized mutual information is

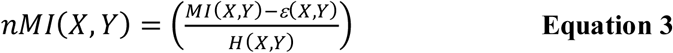

where X and Y are the relevant residue columns. *MI*(*X,Y*) is the mutual information, *H*(*X, Y*) is the joint entropy and *ε*(*X, Y*) is the finite size error.

The mutual information (MI) is given by:

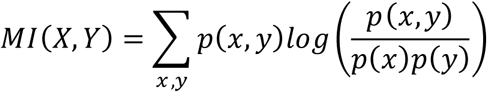

The joint entropy (H) is defined as:

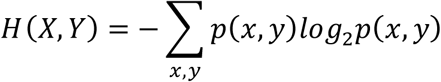

The *sklearn* librarywas used to compute the dynamic couplings between residue sidechain rotation angles and bond distances.

The eigenvector centrality of the C^α^ backbone of the Mg^2+^-free apo-CheY (3CHY.pdb) was characterized earlier based on a structural alphabet that reported secondary structure fluctuations (Wheatley et al., 2020). Here, we used torsional angle as the network metric for C^α^ backbone dynamics (McClendon et al., 2012), The *MI*(*X*(*φ*_1_), *Y*(*φ*_2_)) for the correlations between residue torsional angles (φ, Ψ) was computed as the difference between the conformational self-entropies and the joint entropy.

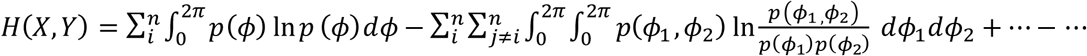

PCA (*gmx anaeig*) for isolation of the principal components and cluster analysis (*gmx cluster*) for determination of conformational heterogeneity complemented the network analysis of the C^α^ backbone in the MD trajectories.

The (*X, Y*) residue states for computation of the dynamic network associated with H-bond transitions were alphanumeric strings that represented distinct H-bond states, similar to the structural alphabets for secondary structure states. By analogy

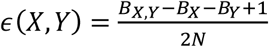

where *N* is sample size, *B*_*X*_, *B*_*Y*_ are the number of distinct residue *X* and *Y* alphanumeric states and *B*_*X,Y*_is their joint probability (see (Carmona et al., 2025)). To give a qualitative illustration, consider the coupling between a R-R_S_ residue R1 with a R-R/W residue R2 that bonds alternately with H_2_O and one residue (not R1). The R1 H-bond will be either OFF or ON, while R2 will have 3 states (OFF, H_2_O ON, residue ON). The nMI will be zero if the 3 R2 states exhibit no preference for whether or not R1 forms a H-bond, and have a maximum value 1, if one R2 state occurs only when the R1 H-bond is ON.

We used betweenness centrality rather than eigenvector centrality for a direct comparison of the fold conformational landscape with the H-bond dynamics. Both centrality measures were defined earlier by (Wheatley et al., 2020) and (Hamid et al., 2022) respectively. In brief, eigenvector centrality taken as the eigenvector of the adjacency matrix measures the connectedness of a node. Betweenness centrality of a node is proportional to the fraction of the shortest paths between other nodes that include it (Rodrigues, 2019). Construction of the covariance matrix (*gmx covar*) preceded the determination of the dynamic couplings and centrality measures. These metrics measured the modulation of the long-range allosteric network by the solvent water at single residue resolution that, in concert with the physical H-bond network compared the allosteric network architecture in the RR superfamily. The betweenness centrality rather than eigenvector centrality was used for a direct comparison of the fold conformational landscape with the H-bond dynamics. The eigenvector centrality taken as the eigenvector of the adjacency matrix measures the connectedness of a node. Betweenness centrality of a node is proportional to the fraction of the shortest paths between other nodes that include it (Rodrigues, 2019). Construction of the covariance matrix (*gmx covar*) preceded the determination of the dynamic couplings and centrality measures.

### Quantitative and statistical analysis

Linear and non-linear best fits to the data, with standard errors, were computed in MATLAB. Pearson correlation and one-way ANOVA were used to compare distributions from replicates within and between different water populations and protein conformations. Mutual information scores were normalized to correct for finite-size error. Decoy distributions generated with custom Python scripts utilizing NumPy and panda libraries assessed score significance. Residue interaction states were randomly shuffled across 1000 iterations to generate a null distribution. The significance evaluation for the DALI distance matrices and the FrustratometerR decoy distributions is detailed in the citations (Holm and Sander, 1993; Parra et al., 2016).

### Figure Preparation

Atomic structure images and movies were prepared in PyMol Chemical bonds were shown with ChemDraw 3D. Cytoscape was used for the 2D representations of the H-bond networks.

## Author Contributions

M.H. Mined databases, analyzed sequence alignments and MD trajectories, wrote custom scripts, ran MD simulations on the LUMS-BIRL cluster, made the figures and prepared the manuscript. I.J.Supervised M.H. advised on MD simulations and edited the manuscript. S.U.C. Supervised M.H., oversaw the LUMS-BIRL computer cluster and prepared the manuscript. S.K. Designed and oversaw the project, ran the Biowulf HPC simulations and wrote the manuscript.

## Acknowledgements

We acknowledge Dr Corie Y. Ralston (C.Y.R) for support, and Drs. Alessandro Pandini and Ann Stock for comments on the manuscript. M.H. was supported by the LUMS Lifesciences graduate program and the Pakistan Higher Education (HEC) grants (20-3629/NRPU/R&D/HEC/14/585**) and** (RF-NCBC-015) (S.U.C). The project was funded by DOE Molecular Foundry grant MF-9084 (S.K/C.Y.R). It utilized the NIH HPC Biowulf computational cluster (*http:// hpc. nih. gov*) supported by the NIH/NINDS intramural program (S.K), the NUST/SINES supercomputing facility (see https://sines.nust.edu.pk for support) (I.J); and the BIRL-LUMS computer cluster (S.U.C). See https://birl.lums.edu.pk */* for BIRL-LUMS cluster support in addition to (20-3629/NRPU/R&D/HEC/14/585), (RF-NCBC-015) .

## Data Availability Statement

The raw MD trajectories have been deposited on Figshare (https://figshare.com/).

### Abbreviations

E. coli / S. typhimurium CheY: (CheY_es_)
Response regulator: (RR)
Molecular dynamics: (MD)
Mutual information: (MI)
Pearson correlation: (P_cc_)
Solvent accessible surface area: (SASA).

## References

1. Attrill, H., P. Gaudet, R.P. Huntley, R.C. Lovering, S.R. Engel, S. Poux, K.M. Van Auken, G. Georghiou, M.C. Chibucos, T.Z. Berardini, V. Wood, H. Drabkin, P. Fey, P. Garmiri, M.A. Harris, T. Sawford, L. Reiser, R. Tauber, S. Toro, and C. Gene Ontology. 2019. Annotation of gene product function from high-throughput studies using the Gene Ontology. Database (Oxford). 2019.

2. Biswas, M., S. Dey, S. Khamrui, U. Sen, and J. Dasgupta. 2013. Conformational barrier of CheY3 and inability of CheY4 to bind FliM control the flagellar motor action in Vibrio cholerae. PLoS One. 8:e73923.

3. Bondar, A.N. 2022. Graphs of Hydrogen-Bond Networks to Dissect Protein Conformational Dynamics. J Phys Chem B. 126:3973–3984.

4. Bordin, N., H. Scholes, C. Rauer, J. Roca-Martinez, I. Sillitoe, and C. Orengo. 2024. Clustering protein functional families at large scale with hierarchical approaches. Protein Sci. 33:e5140.

5. Bordin, N., I. Sillitoe, V. Nallapareddy, C. Rauer, S.D. Lam, V.P. Waman, N. Sen, M. Heinzinger, M. Littmann, S. Kim, S. Velankar, M. Steinegger, B. Rost, and C. Orengo. 2023. AlphaFold2 reveals commonalities and novelties in protein structure space for 21 model organisms. Commun Biol. 6:160.

6. Bourret, R.B., J.F. Hess, and M.I. Simon. 1990. Conserved aspartate residues and phosphorylation in signal transduction by the chemotaxis protein CheY. Proc Natl Acad Sci U S A. 87:41–45.

7. Carmona, O.G., J. Kleinjung, D. Anastasiou, C. Oostenbrink, and F. Fraternali. 2025. AllohubPy: Detecting Allosteric Signals Through An Information-theoretic Approach. J Mol Biol:168969.

8. Cho, H.S., S.Y. Lee, D. Yan, X. Pan, J.S. Parkinson, S. Kustu, D.E. Wemmer, and J.G. Pelton. 2000. NMR structure of activated CheY. J Mol Biol. 297:543–551.

9. Das, S., D. Lee, I. Sillitoe, N.L. Dawson, J.G. Lees, and C.A. Orengo. 2015. Functional classification of CATH superfamilies: a domain-based approach for protein function annotation. Bioinformatics. 31:3460–3467.

10. Dyer, C.M., and F.W. Dahlquist. 2006. Switched or not?: the structure of unphosphorylated CheY bound to the N terminus of FliM. J Bacteriol. 188:7354–7363.

11. Dyer, C.M., M.L. Quillin, A. Campos, J. Lu, M.M. McEvoy, A.C. Hausrath, E.M. Westbrook, P. Matsumura, B.W. Matthews, and F.W. Dahlquist. 2004. Structure of the constitutively active double mutant CheYD13K Y106W alone and in complex with a FliM peptide. J Mol Biol. 342:1325–1335.

12. Ferreiro, D.U., J.A. Hegler, E.A. Komives, and P.G. Wolynes. 2007. Localizing frustration in native proteins and protein assemblies. Proc Natl Acad Sci U S A. 104:19819–19824.

13. Fersht, A.R., J.P. Shi, J. Knill-Jones, D.M. Lowe, A.J. Wilkinson, D.M. Blow, P. Brick, P. Carter, M.M. Waye, and G. Winter. 1985. Hydrogen bonding and biological specificity analysed by protein engineering. Nature. 314:235–238.

14. Foster, C.A., R.E. Silversmith, R.M. Immormino, L.R. Vass, E.N. Kennedy, Y. Pazy, E.J. Collins, and R.B. Bourret. 2021. Role of Position K+4 in the Phosphorylation and Dephosphorylation Reaction Kinetics of the CheY Response Regulator. Biochemistry. 60:2130–2151.

15. Foster, C.A., and A.H. West. 2017. Use of restrained molecular dynamics to predict the conformations of phosphorylated receiver domains in two-component signaling systems. Proteins. 85:155–176.

16. Fraiberg, M., O. Afanzar, C.K. Cassidy, A. Gabashvili, K. Schulten, Y. Levin, and M. Eisenbach. 2015. CheY’s acetylation sites responsible for generating clockwise flagellar rotation in Escherichia coli. Mol Microbiol. 95:231–244.

17. Franken, G.A.C., M.A. Huynen, L.A. Martinez-Cruz, R.J.M. Bindels, and J.H.F. de Baaij. 2022. Structural and functional comparison of magnesium transporters throughout evolution. Cell Mol Life Sci. 79:418.

18. Freiberger, M.I., V. Ruiz-Serra, C. Pontes, M. Romero-Durana, P. Galaz-Davison, C.A. Ramirez-Sarmiento, C.D. Schuster, M.A. Marti, P.G. Wolynes, D.U. Ferreiro, R.G. Parra, and A. Valencia. 2023. Local energetic frustration conservation in protein families and superfamilies. Nat Commun. 14:8379.

19. Fu, L., B. Niu, Z. Zhu, S. Wu, and W. Li. 2012. CD-HIT: accelerated for clustering the next-generation sequencing data. Bioinformatics. 28:3150–3152.

20. Ganguli, S., H. Wang, P. Matsumura, and K. Volz. 1995. Uncoupled phosphorylation and activation in bacterial chemotaxis. The 2.1-A structure of a threonine to isoleucine mutant at position 87 of CheY. J Biol Chem. 270:17386–17393.

21. Gao, R., S. Bouillet, and A.M. Stock. 2019. Structural Basis of Response Regulator Function. Annu Rev Microbiol. 73:175–197.

22. Gao, R., T.R. Mack, and A.M. Stock. 2007. Bacterial response regulators: versatile regulatory strategies from common domains. Trends Biochem Sci. 32:225–234.

23. Giannetti, G., F. Matsumura, F. Caporaletti, D. Micha, G.H. Koenderink, I.M. Ilie, M. Bonn, S. Woutersen, and G. Giubertoni. 2025. Water and Collagen: A Mystery Yet to Unfold. Biomacromolecules. 26:2784–2799.

24. Hamid, M., S.U. Chaudhary, A. Pandini, and S. Khan. 2025. Allosteric network analysis toolkit for single domain phosphoproteins. Methods in Molecular Biology. Protein Design & Evolution.

25. Hamid, M., M.F. Khalid, S.U. Chaudhary, and S. Khan. 2022. The Solvation of the E. coli CheY Phosphorylation Site Mapped by XFMS. Int J Mol Sci. 23.

26. Holm, L. 2022. Dali server: structural unification of protein families. Nucleic Acids Res. 50:W210–W215.

27. Holm, L., A. Laiho, P. Toronen, and M. Salgado. 2023. DALI shines a light on remote homologs: One hundred discoveries. Protein Sci. 32:e4519.

28. Holm, L., and C. Sander. 1993. Protein structure comparison by alignment of distance matrices. J Mol Biol. 233:123–138.

29. Holm, L., and C. Sander. 1995. Dali: a network tool for protein structure comparison. Trends Biochem Sci. 20:478–480.

30. Immormino, R.M., R.E. Silversmith, and R.B. Bourret. 2016. A Variable Active Site Residue Influences the Kinetics of Response Regulator Phosphorylation and Dephosphorylation. Biochemistry. 55:5595–5609.

31. Jiang, M., R.B. Bourret, M.I. Simon, and K. Volz. 1997. Uncoupled phosphorylation and activation in bacterial chemotaxis. The 2.3 A structure of an aspartate to lysine mutant at position 13 of CheY. J Biol Chem. 272:11850–11855.

32. Jumper, J., R. Evans, A. Pritzel, T. Green, M. Figurnov, O. Ronneberger, K. Tunyasuvunakool, R. Bates, A. Zidek, A. Potapenko, A. Bridgland, C. Meyer, S.A.A. Kohl, A.J. Ballard, A. Cowie, B. Romera-Paredes, S. Nikolov, R. Jain, J. Adler, T. Back, S. Petersen, D. Reiman, E. Clancy, M. Zielinski, M. Steinegger, M. Pacholska, T. Berghammer, S. Bodenstein, D. Silver, O. Vinyals, A.W. Senior, K. Kavukcuoglu, P. Kohli, and D. Hassabis. 2021. Highly accurate protein structure prediction with AlphaFold. Nature. 596:583–589.

33. Kadaoluwa Pathirannahalage, S.P., N. Meftahi, A. Elbourne, A.C.G. Weiss, C.F. McConville, A. Padua, D.A. Winkler, M. Costa Gomes, T.L. Greaves, T.C. Le, Q.A. Besford, and A.J. Christofferson. 2021. Systematic Comparison of the Structural and Dynamic Properties of Commonly Used Water Models for Molecular Dynamics Simulations. J Chem Inf Model. 61:4521–4536.

34. Karplus, K., K. Sjolander, C. Barrett, M. Cline, D. Haussler, R. Hughey, L. Holm, and C. Sander. 1997. Predicting protein structure using hidden Markov models. Proteins. Suppl 1:134–139.

35. Kennedy, E.N., C.A. Foster, S.A. Barr, and R.B. Bourret. 2022. General strategies for using amino acid sequence data to guide biochemical investigation of protein function. Biochem Soc Trans. 50:1847–1858.

36. Khan, S. 2022. Conformational spread drives the evolution of the calcium-calmodulin protein kinase II. Sci Rep. 12:8499.

37. Khan, S., J.L. Spudich, J.A. McCray, and D.R. Trentham. 1995. Chemotactic signal integration in bacteria. Proc Natl Acad Sci U S A. 92:9757–9761.

38. Khemaissa, S., A. Walrant, and S. Sagan. 2022. Tryptophan, more than just an interfacial amino acid in the membrane activity of cationic cell-penetrating and antimicrobial peptides. Q Rev Biophys. 55:e10.

39. Lee, S.Y., H.S. Cho, J.G. Pelton, D. Yan, E.A. Berry, and D.E. Wemmer. 2001. Crystal structure of activated CheY. Comparison with other activated receiver domains. J Biol Chem. 276:16425–16431.

40. Li, W., and A. Godzik. 2006. Cd-hit: a fast program for clustering and comparing large sets of protein or nucleotide sequences. Bioinformatics. 22:1658–1659.

41. Lukat, G.S., B.H. Lee, J.M. Mottonen, A.M. Stock, and J.B. Stock. 1991. Roles of the highly conserved aspartate and lysine residues in the response regulator of bacterial chemotaxis. J Biol Chem. 266:8348–8354.

42. Lukat, G.S., W.R. McCleary, A.M. Stock, and J.B. Stock. 1992. Phosphorylation of bacterial response regulator proteins by low molecular weight phospho-donors. Proc Natl Acad Sci U S A. 89:718–722.

43. Margreitter, C., M.M. Reif, and C. Oostenbrink. 2017. Update on phosphate and charged post-translationally modified amino acid parameters in the GROMOS force field. J Comput Chem. 38:714–720.

44. Mark, P., and L. Nilsson. 2002. Structure and dynamics of liquid water with different long-range interaction truncation and temperature controlmethods in moleculardynamics simulations. J Comput Chem. 23:1211–1219.

45. McClendon, C.L., L. Hua, A. Barreiro, and M.P. Jacobson. 2012. Comparing Conformational Ensembles Using the Kullback-Leibler Divergence Expansion. J Chem Theory Comput. 8:2115–2126.

46. Nakamura, T., K.D. Yamada, K. Tomii, and K. Katoh. 2018. Parallelization of MAFFT for large-scale multiple sequence alignments. Bioinformatics. 34:2490–2492.

47. Pace, C.N., G. Horn, E.J. Hebert, J. Bechert, K. Shaw, L. Urbanikova, J.M. Scholtz, and J. Sevcik. 2001. Tyrosine hydrogen bonds make a large contribution to protein stability. J Mol Biol. 312:393–404.

48. Page, S.C., R.M. Immormino, T.H. Miller, and R.B. Bourret. 2016. Experimental Analysis of Functional Variation within Protein Families: Receiver Domain Autodephosphorylation Kinetics. J Bacteriol. 198:2483–2493.

49. Pandini, A., J. Kleinjung, S. Rasool, and S. Khan. 2015. Coevolved Mutations Reveal Distinct Architectures for Two Core Proteins in the Bacterial Flagellar Motor. PLoS One. 10:e0142407.

50. Papoian, G.A., J. Ulander, M.P. Eastwood, Z. Luthey-Schulten, and P.G. Wolynes. 2004. Water in protein structure prediction. Proc Natl Acad Sci U S A. 101:3352–3357.

51. Parra, R.G., M.I. Freiberger, M. Poley-Gil, M. Fernandez-Martin, L.G. Radusky, V. Ruiz-Serra, P.G. Wolynes, D.U. Ferreiro, and A. Valencia. 2024. Frustraevo: a web server to localize and quantify the conservation of local energetic frustration in protein families. Nucleic Acids Res. 52:W233–W237.

52. Parra, R.G., N.P. Schafer, L.G. Radusky, M.Y. Tsai, A.B. Guzovsky, P.G. Wolynes, and D.U. Ferreiro. 2016. Protein Frustratometer 2: a tool to localize energetic frustration in protein molecules, now with electrostatics. Nucleic Acids Res. 44:W356–360.

53. Pavlopoulos, G.A., F.A. Baltoumas, S. Liu, O. Selvitopi, A.P. Camargo, S. Nayfach, A. Azad, S. Roux, L. Call, N.N. Ivanova, I.M. Chen, D. Paez-Espino, E. Karatzas, C. Novel Metagenome Protein Families, I. Iliopoulos, K. Konstantinidis, J.M. Tiedje, J. Pett-Ridge, D. Baker, A. Visel, C.A. Ouzounis, S. Ovchinnikov, A. Buluc, and N.C. Kyrpides. 2023. Unraveling the functional dark matter through global metagenomics. Nature. 622:594–602.

54. Pazy, Y., A.C. Wollish, S.A. Thomas, P.J. Miller, E.J. Collins, R.B. Bourret, and R.E. Silversmith. 2009. Matching biochemical reaction kinetics to the timescales of life: structural determinants that influence the autodephosphorylation rate of response regulator proteins. J Mol Biol. 392:1205–1220.

55. Petsko, G.A. 2000. Chemistry and biology. Proc Natl Acad Sci U S A. 97:538–540.

56. Prisant, M.G., C.J. Williams, V.B. Chen, J.S. Richardson, and D.C. Richardson. 2020. New tools in MolProbity validation: CaBLAM for CryoEM backbone, UnDowser to rethink "waters," and NGL Viewer to recapture online 3D graphics. Protein Sci. 29:315–329.

57. Rodrigues, F.A. 2019. Network centrality: An introduction. arXiv.

58. Rogacheva, O.N., S.A. Izmailov, L.V. Slipchenko, and N.R. Skrynnikov. 2017. A new structural arrangement in proteins involving lysine NH(3)(+) group and carbonyl. Sci Rep. 7:16402.

59. Rojano, E., F.M. Jabato, J.R. Perkins, J. Cordoba-Caballero, F. Garcia-Criado, I. Sillitoe, C. Orengo, J.A.G. Ranea, and P. Seoane-Zonjic. 2022. Assigning protein function from domain-function associations using DomFun. BMC Bioinformatics. 23:43.

60. Roman, S.J., M. Meyers, K. Volz, and P. Matsumura. 1992. A chemotactic signaling surface on CheY defined by suppressors of flagellar switch mutations. J Bacteriol. 174:6247–6255.

61. Samways, M.L., R.D. Taylor, H.E. Bruce Macdonald, and J.W. Essex. 2021. Water molecules at protein-drug interfaces: computational prediction and analysis methods. Chem Soc Rev. 50:9104–9120.

62. Schiebel, J., R. Gaspari, T. Wulsdorf, K. Ngo, C. Sohn, T.E. Schrader, A. Cavalli, A. Ostermann, A. Heine, and G. Klebe. 2018. Intriguing role of water in protein-ligand binding studied by neutron crystallography on trypsin complexes. Nat Commun. 9:3559.

63. Siemers, M., and A.N. Bondar. 2021. Interactive Interface for Graph-Based Analyses of Dynamic H-Bond Networks: Application to Spike Protein S. J Chem Inf Model.

64. Siemers, M., M. Lazaratos, K. Karathanou, F. Guerra, L.S. Brown, and A.N. Bondar. 2019. Bridge: A Graph-Based Algorithm to Analyze Dynamic H-Bond Networks in Membrane Proteins. J Chem Theory Comput. 15:6781–6798.

65. Sillitoe, I., N. Bordin, N. Dawson, V.P. Waman, P. Ashford, H.M. Scholes, C.S.M. Pang, L. Woodridge, C. Rauer, N. Sen, M. Abbasian, S. Le Cornu, S.D. Lam, K. Berka, I.H. Varekova, R. Svobodova, J. Lees, and C.A. Orengo. 2021. CATH: increased structural coverage of functional space. Nucleic Acids Res. 49:D266–D273.

66. Smith, S.O., M. Eilers, D. Song, E. Crocker, W. Ying, M. Groesbeek, G. Metz, M. Ziliox, and S. Aimoto. 2002. Implications of threonine hydrogen bonding in the glycophorin A transmembrane helix dimer. Biophys J. 82:2476–2486.

67. Spyrakis, F., M.H. Ahmed, A.S. Bayden, P. Cozzini, A. Mozzarelli, and G.E. Kellogg. 2017. The Roles of Water in the Protein Matrix: A Largely Untapped Resource for Drug Discovery. J Med Chem. 60:6781–6827.

68. Stock, A.M., E. Martinez-Hackert, B.F. Rasmussen, A.H. West, J.B. Stock, D. Ringe, and G.A. Petsko. 1993. Structure of the Mg(2+)-bound form of CheY and mechanism of phosphoryl transfer in bacterial chemotaxis. Biochemistry. 32:13375–13380.

69. Stock, A.M., J.M. Mottonen, J.B. Stock, and C.E. Schutt. 1989. Three-dimensional structure of CheY, the response regulator of bacterial chemotaxis. Nature. 337:745–749.

70. Straughn, P.B., L.R. Vass, C. Yuan, E.N. Kennedy, C.A. Foster, and R.B. Bourret. 2020. Modulation of Response Regulator CheY Reaction Kinetics by Two Variable Residues That Affect Conformation. J Bacteriol. 202.

71. Tani, K., and Y. Fujiyoshi. 2014. Water channel structures analysed by electron crystallography. Biochim Biophys Acta. 1840:1605–1613.

72. Toro-Roman, A., T.R. Mack, and A.M. Stock. 2005. Structural analysis and solution studies of the activated regulatory domain of the response regulator ArcA: a symmetric dimer mediated by the alpha4-beta5-alpha5 face. J Mol Biol. 349:11–26.

73. Tsai, C.S., and S.C. Winans. 2010. LuxR-type quorum-sensing regulators that are detached from common scents. Mol Microbiol. 77:1072–1082.

74. Volz, K., and P. Matsumura. 1991. Crystal structure of Escherichia coli CheY refined at 1.7-A resolution. J Biol Chem. 266:15511–15519.

75. Wheatley, P., S. Gupta, Y. Chen, C.J. Petzold, C.R. Ralston, D.F. Blair, and S. Khan. 2020. Allosteric priming of E. coli CheY by the flagellar motor protein FliM. Biophysical Journal:1108–1122.

76. Zhu, X., J. Rebello, P. Matsumura, and K. Volz. 1997a. Crystal structures of CheY mutants Y106W and T87I/Y106W. CheY activation correlates with movement of residue 106. *J Biol Chem*. 272:5000-5006.

77. Zhu, X., K. Volz, and P. Matsumura. 1997b. The CheZ-binding surface of CheY overlaps the CheA-and FliM-binding surfaces. J Biol Chem. 272:23758–23764.

