## Supplementary Material for "Conservation of the hydrogen-bond network in bacterial response regulators"

*Submitted BioRxiv*

### FIGURES

**Figure S1.** Principal component analysis of CheY conformational landscapes.

**Figure S2.** Structural determinants of water mobility.

**Figure S3.** nMI coupling strength versus state type and number.

### VIDEOS

**Video S1.** Local frustration in the RR superfamily.

**Video S2.** 2CHE hydrogen-bonded network.

**Video S3.** $1FQW_{PO4}$ hydrogen-bonded network.

### REPOSITORY

Raw trajectory files have been deposited on (*https://figshare.com/*). They will be made public upon publication.

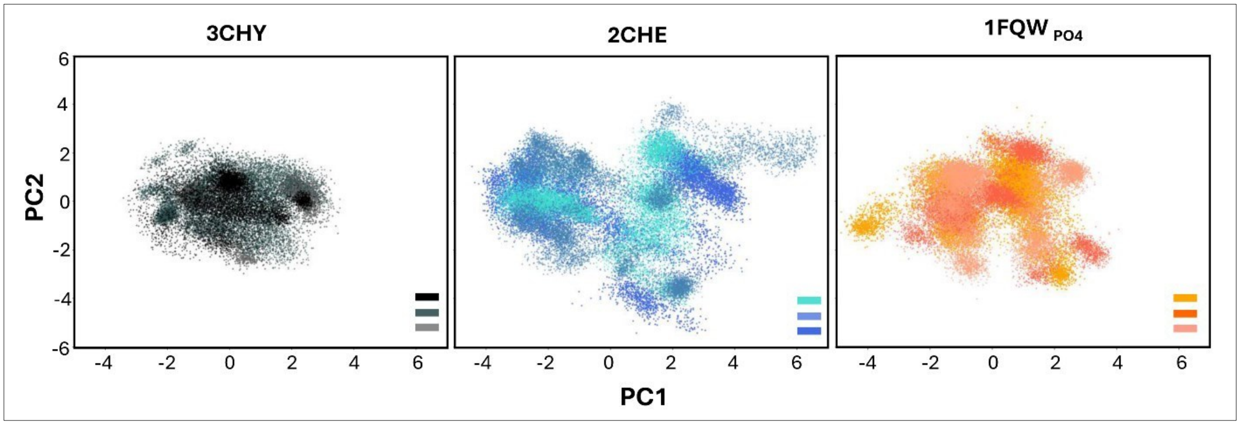


***Figure S1: Principal component analysis of CheY conformational landscapes.*** *Variations between replicate (n = 3) trajectories obtained for the three different CheY forms – apo (3CHY), basal, $Mg^{2+}$-bound.(2CHE)) and phosphorylated, $Mg^{2+}$-bound ($1FQW_{PO4}$). The 3CHY replicates were 0.6, 0.63 and 1 μs. The 2CHE and $1FQW_{PO4}$ replicates were 1 μs each. Different shades (horizontal bars) represent different replicates. Comparison of the 3CHY and 2CHE panels suggests that, after correction for trajectory size, the bound $Mg^{2+}$may increase the accessible conformational landscape. Comparison of the 2CHE and $1FQW_{PO4}$ panels suggests that phosphorylation selects, rather than increase, the basal CheY substates in the landscape.*

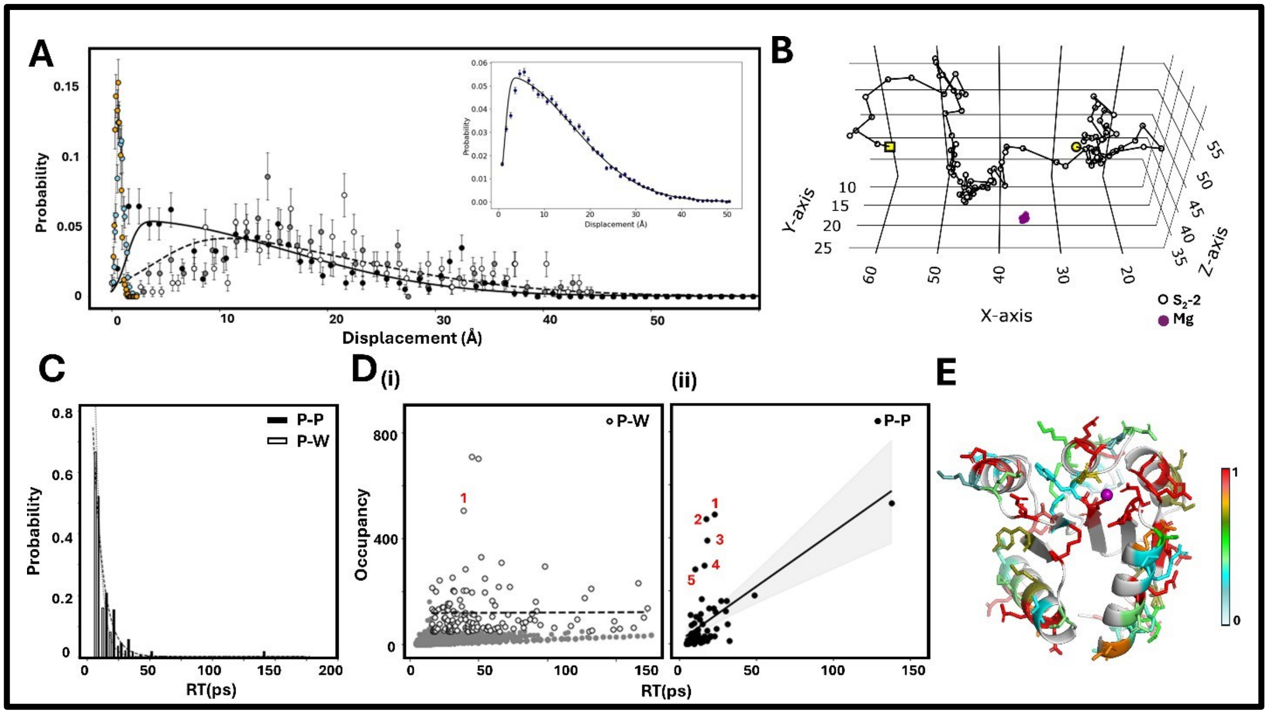


***Figure S3: Structural determinants of water mobility.*** ***A.*** *Mobility of crystal waters plotted as the distribution of the distance (mean +/- se) from the starting coordinates in successive frames over the 500ps period (3 replicates).* $Mg^{2+}$ *coordination: 1st shell (2CHE (cyan, n=3),* $1FQW_{PO4}$ *(orange, n=2). Fit (dotted line)* $\frac{1}{0.3\sqrt{2\pi}}.exp(-\frac{(x-0.756)^2}{2(0.3)^2})$. *Peak distance = 0.76Å, 2nd shell (2CHE) 3 waters (white, gray, black symbols). Radial distances of solvation ring =1st Shell (1.96Å ± 0.009 Å), 2nd shell (29.1±1.14Å).. 2nd shell waters reset to crystal water positions (4.5±0.7Å) for production runs. Skewed Gaussian fits (dashed line* $(2*\phi(x)*\Phi(4.4*x)$; *solid line* $2*\phi(x)*\Phi(16.0*x)$, *where x =distance,* $\phi$ *= probability density function,* $\Phi$ *= cumulative distribution function.* ***Inset*** *shows fit to "other" crystal waters. Compare with solid line 2nd shell water fit.* ***B****. Example trajectory of a 2nd shell water (yellow symbols mark start (circle) and end (square)).* $Mg^{2+}$ *(magenta) displacement over 500 ps.* ***C.*** *Residence time (RT) distributions for P-P and P-W H-bonds (3 replicates, 4 ns duration, 5 ps intervals).* $Mg^{2+}$ *coordination components excluded.* ***D.*** *The correlation between H-bond occupancy(Occ) and residence time (RT). Occupancy filter = 0.05* ***(i)*** *P-W bonds. H-bonds that remain (open symbols) or are eliminated (grey symbols) by the occupancy filter. Fit (dashed line) Occ=0.02(RT)+118.1. Non-crystal water trapped at 2nd water shell location (asterisk). (ii) P-P bonds. Fit (solid line with 95% confidence region shaded gray) Occ= 4.09(RT)+9.7.* ***P-W.*** *D12-*$HOH_{3948}$*(1)).* $HOH_{3948}$ *acts as a 2nd shell water for a substantial fraction of one 4 ns trajectory, with 6.03±0.06Å distance from the* $Mg^{2+}$ *ion over the final 3.1 ns.* ***P-P*** *(R18-E35(1), D41-K45(2), T112-T115(3), M17-K109(4), R22-E35(5)).* ***E****. Water (W)contact map after 1st persistence filter (> 3 frames). Residues that form P-W bonds (stick sidechains) are color coded based by their P-W occupancy (Bar). Acidic (D,E) residues had the highest occupancy(0.95±0.01 (n=18)), followed by basic (K,R) residues (0.54±0.6 (n=15)). T87 occupancy was notably higher (0.83) than other threonine residues (0.43±0.11(n=5)). Two aromatic residues (W58 (0.20), Y106 (0.61)) also formed P-W bonds. Methionine has the lowest*

*occupancies (0.10±0.04 (n=6)). Other polar residues (N,S,Q) made up the remaining population (0.56±0.05 (n=15)) There was no correlation with the SASA ($P_{CC}$= -0.16). 4 ns 2CHE trajectory.*

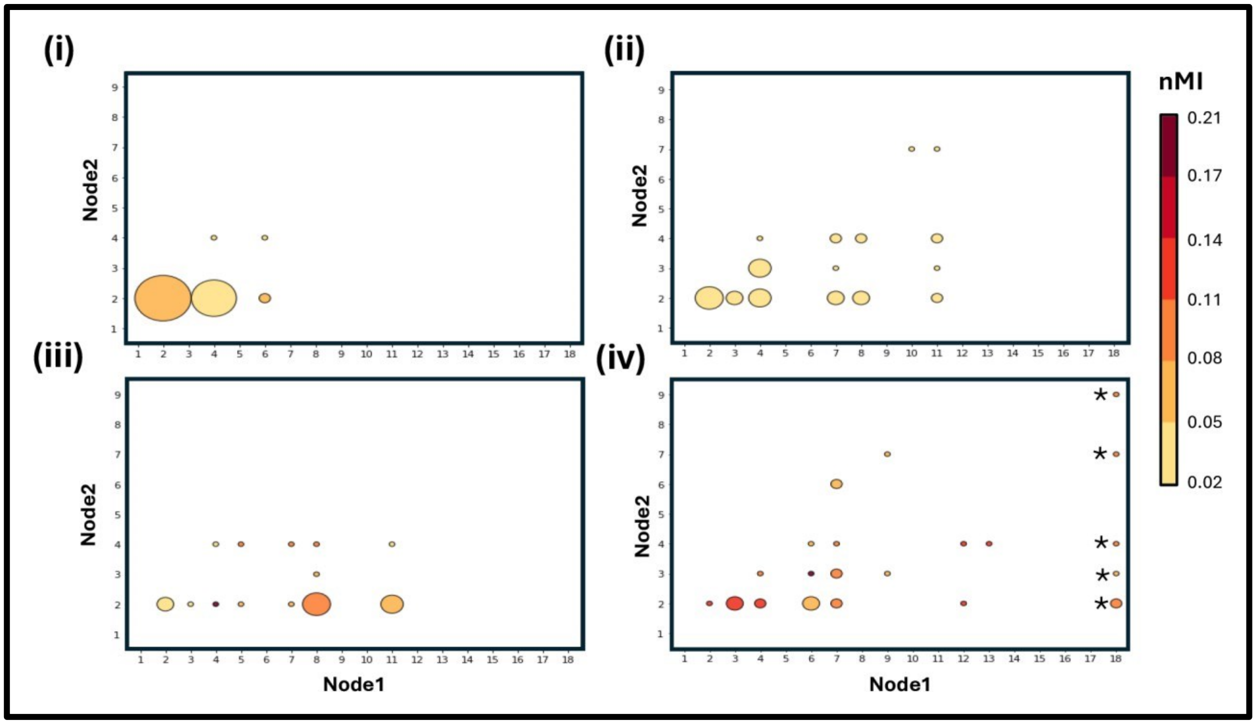


***Figure S3: State multiplicity for the dynamic couplings. (i)*** *2CHE.* ***(ii)*** *1FQW$_{PO4}$.* ***(iii)*** *4H6O.* ***(iv)*** *4H6O$_{PO4}$. Significant couplings were identified by comparison with a decoy distribution. Their state multiplicity is shown with the state counts of the residue node with the higher count per coupling on the x-axis, and the lower count node on the y-axis. Symbol size represents the number of independent couplings and color the mean NMI value indicated by the bar. 3x20 ns trajectories. D57$_{PO4}$ couplings (asterisks (*)). 3 20ns replicates*
